# Peripheral nerve derived extracellular vesicles are dynamically regulated in chemotherapy induced painful neuropathy

**DOI:** 10.64898/2026.08.20.746051

**Authors:** Miles Vecchitto, Gail Funk, Zixuan Wang, Takahito Arai, Stefano Martellucci, Saptarshi Sinha, Aaron Tran, Masaki Norimoto, Majid Ghassamian, Pradipta Ghosh, Steven L. Gonias, Wendy M. Campana

**Author notes:** To who correspondence should be addressed: Wendy M Campana, PhD.

## Abstract

Peripheral neuropathy is typically associated with neuropathic pain, which arises from nerve damage and altered communication between peripheral glia and sensory axons. Although extracellular vesicles (EVs) are critical mediators of intercellular communication, their role in healthy and neuropathic peripheral nerves remains poorly understood. We developed a minimally disruptive protocol to isolate sciatic nerve-derived EVs (snEVs) under healthy and neuropathic conditions. While snEVs are inherently heterogeneous, LC-MS/MS proteomic profiling revealed that healthy snEVs are selectively enriched in proteins essential for axonal integrity and Schwann cell (SC) myelination. In contrast, snEVs isolated from mice treated with paclitaxel (PTX), a cancer chemotherapy agent known to cause neuropathic pain, displayed a markedly distinct proteomic signature, uniquely enriched in metabolic enzymes and receptors linked to neuronal excitability. Functionally, neuropathic snEVs acutely activated pain-associated intracellular cell-signaling pathways in sensory neurons. To determine whether painful peripheral neuropathy (PPN) alters SC-derived snEV subpopulations *in vivo*, we generated a novel P0-Cre-turbo-GFP/human-CD9-EV reporter mouse model. High-resolution microscopy and single fluorescence nanoparticle tracking analysis revealed that nerves from mice treated with PTX yielded fewer, yet significantly larger, SC-derived snEVs. These findings demonstrate that peripheral nerve EV dynamics and cargo are actively remodeled in PPN.

## Introduction

Painful peripheral neuropathy (PPN) is a common syndrome emerging from insult to a peripheral nerve that may be metabolic, toxic or traumatic. Chemotherapy-induced painful peripheral neuropathy (CIPPN) results from cancer treatment and is highly prevalent in cancer patients (Serenty et al., 2014). Paclitaxel (PTX), one of the most effective and commonly used chemotherapy agents, is dose-limiting due to CIPPN that persists long after treatment is complete (Maiaru et al., 2025). PTX-induced neuropathic pain is associated with sciatic nerve dysfunction and metabolic imbalances in Schwann cells (SCs), the major glia in peripheral nerves. In myelinated SCs, PTX damages the myelin sheath and reduces expression of compact myelin proteins such as P0. However, the SC-specific mechanisms underlying CIPPN remain unclear. Currently, there are no effective treatments to prevent or limit development of CIPPN. Thus, a better understanding of cellular and molecular mechanism is essential.

Extracellular vesicles (EVs) are lipid bilayer-enclosed particles produced ubiquitously by eukaryotic cells (Jeppesen *et al*., 2019; van Niel *et al*., 2018). By transferring macromolecules from cells of origin to target cells, EVs are highly involved in cell-cell communication that is essential in health and disease (Sato and Weaver, 2018; Gurung *et al*. 2021). In addition to their ability to transfer cargo (proteins, RNA, DNA, microRNA) from the cell of origin to target cells (Lässer *et al*., 2018), EVs also regulate cell physiology through interactions between EV surface proteins and receptors on the target cell (Gonias *et al*., 2022). It has been proposed that the substantial membrane surface of an EV may present multiple ligands and thereby cluster target cell receptors to induce EV-dependent cell-signaling (Jahnke and Staufer, 2024). EVs are abundant in body fluids including blood (Iannotta *et al*., 2024) and milk (Leiferman *et al*., 2019) and participate in autocrine, paracrine, and endocrine cell-cell communication. However, challenges remain in understanding the activities of EVs *in vivo*. Tissue-derived EVs have the potential to provide a molecular signature of the local tissue microenvironment (Lee *et al*., 2024) which cannot be replicated when EVs are collected from cell culture conditioned medium.

After peripheral injury, SC-derived EVs have been shown to activate regenerative programs by transferring to axons functional molecular cargo that promotes axonal elongation (Lopez-Verrilli *et al*., 2013; Court and Alvarez, 2016; López-Leal *et al*., 2020; Marín-Venegas and Court, 2026; Ching *et al*., 2018). SC-derived EVs also possess the ability to dampen inflammation, exemplified by decoy TNFR1-bearing EVs that sequester excess TNFα (Sadri *et al*., 2022). However, studies in peripheral nerve regeneration and neuropathic pain, performed to date, have mostly utilized EVs from cultured SCs, which may not accurately reflect EVs generated *in vivo* in intact nerves. Whether pathological shifts in peripheral nerve EV subpopulations, cargo, and paracrine signaling contribute to the initiation or maintenance of neuropathic pain remains a fundamental yet unresolved question.

Here, we established a novel method for isolating tissue-derived EVs from mouse sciatic nerves (snEVs) without compromising cellular integrity. To determine how healthy snEVs differ from those associated with CIPPN neuropathic pain, we compared the proteomes of both phenotypes. Neuropathic snEVs exhibited a distinct protein signature linked to neuronal excitability and activated pain-associated cell signaling in primary sensory neuron cultures. We then generated a novel inducible P0-Cre-tGFP-EV reporter mouse, which expresses a tGFP-CD9 fusion protein specifically in Schwann cells (SCs), to track SC-derived snEVs during CIPPN *in vivo*. SC-derived snEVs were significantly fewer and larger in mice exhibiting pain-related behavior. Together, these findings suggest that changes in snEV protein cargo and altered SC-derived snEV populations play an instrumental role in the response to nerve damage during painful peripheral neuropathy.

## MATERIALS and METHODS

### Ethics statement

All animal housing, breeding, and care protocols were performed with approval by the American Association for Assessment and Accreditation of Laboratory Animal Care (AAALAC). All animal experimentation was approved by the Institutional Animal Care and Use Committee (IACUC) at the University of California, San Diego (UCSD) and the University of California, Davis (UCD). The Veterans Administration (VA) San Diego Healthcare System also approved animal studies in accordance with the Guidelines for Use of Animals in Research through U.S. Department of Agriculture Animal Welfare Act (AWA) regulations and the National Institute of Health’s (NIH) Public Health Service policy (PHS).

### Animals

C57BL/6J mice and transgenic inducible turbo GFP/human CD9 (tGFP/hCD9) EV reporter (TIGER) mice (B6;129S1-Gt(ROSA)^26Sortm1(CAG-CD9/GFP)Dmfel/J^) were obtained from Jackson Laboratory (033361, 000664). TIGER mice were crossed with P0-Cre mice in the BL6/C57 background originally obtained from Dr. Laura Feltri (Feltri *et al*., 1999) and published previously (Orita *et al*., 2013; Martellucci et al., 2025). These mice express Cre recombinase under the control of the myelin protein zero (P0) promoter. Progeny that were hemizygous for *P0-Cre* were bred with TIGER mice to generate experimental animals (PT mice). Approximately 25% of the resulting pups were homozygous for tGFP and hemizygous for *Cre*, following Mendelian genetics. Cre-negative male and female littermates were studied as negative controls. All mice were housed using a 12 h:12 h light:dark cycle with *ad libitum* access to food and water.

### Isolation of sciatic nerve-derived EVs

Adult mice (25 g) were anaesthetized using 3-5% isoflurane with an oxygen flow rate of 2 L/minute. Two to 10 sciatic nerves were isolated and placed in Dulbecco’s Modified Eagle Medium (DMEM; Gibco, 11885) supplemented with 25 mM N-2-hydroxyethylpiperazine-N-2-ethane sulfonic acid buffer (HEPES; Gibco, 15630-080) followed by the removal of the epineural sheath. The nerves were then digested in a collagenase/dispase blend (C1D; Sigma Aldrich, SCR139) in DMEM for 1 h at 37°C (Martellucci *et al*., 2024). Digested nerves were gently titurated ten times, using an 18-G (Fisher Scientific, 14-826-5G) and 23-G (Fisher Scientific, 14-826-6B) needle in succession. Intact cells were pelleted gently by centrifugation at 1,500 x *g* (Beckman, Allegra X-30R) for 10 minutes at 4°C. Larger particles were pelleted from the supernatant at 10,000 x *g* for 10 minutes at 4°C. EVs were then extracted from the supernatant using a qEVoriginal 35 nm size exclusion chromatography (SEC) column (Izon, ICO-35; Izon Science) using sterile-filtered (GenClone, 25-240) Dulbecco’s phosphate-buffered saline (DPBS; Gibco, 14190144) as the elution buffer. The addition of the prepared nerve digest was counted as fraction 1 and EVs were found to be isolated in fractions 7-9. CaptoCore (Sigma) was utilized for further purification.

### Immunoblotting

EV or tissue extract samples were loaded onto precast 10% or 12% polyacrylamide gels (Bio-Rad, 4561034 and 4561044, respectively). EV concentration of each sample was determined by NTA (Malvern) and tissue lysate by bicinchoninic acid assay (BCA; Thermo Fisher Scientific, 23226). Samples were subjected to SDS-PAGE and electro-transferred to nitrocellulose membranes as previously described (Martellucci *et al*., 2024). Membranes were blocked with 5% nonfat dry milk (Bio-Rad, 1706404) dissolved in Tris-buffered saline (10 mM Tris-HCl, 150 mM NaCl, pH 7.5) with 0.1% Tween-20 (TBS-T) for 1 hour at room temperature. Primary antibodies were diluted in blocking solution or 5% bovine serum albumin (BSA)/TBS-T (WSR Life Sciences, 0332-25G) and were incubated overnight at 4°C or for 1 hour at room temperature. The following primary antibodies were used: anti-flotillin-1 (rabbit polyclonal 1:1000; Cell Signaling Technologies, 3253S), anti-TSG101 (rabbit monoclonal 1:1000; Abcam, ab125011), anti-CD9 (rabbit monoclonal 1:1000; Abcam, ab307085, rabbit monoclonal 1:1500; Novus, nbp2-67310), anti-myelin protein zero (P0) (rabbit monoclonal 1:1000; Abcam, ab183868), anti-GM130 (rabbit monoclonal 1:1000; Thermo Fisher Scientific, ARC0589), anti-βIII-tubulin (rabbit monoclonal 1:1000; Cell Signaling Technologies, 5666S), anti-β-actin (mouse monoclonal 1:1000; Cell Signaling Technologies, 3700S), anti-annexin-2 (mouse monoclonal 1:1000; BD Biosciences, bd610069), anti-SOX10 (rabbit monoclonal 1:1000; Invitrogen, MA5-32335), anti-calnexin (rabbit polyclonal 1:1000; Enzo Life Sciences, ADI-SPA-860-F), anti-phospho-AKT (Ser473) or (Thr308) (rabbit polyclonal 1:1000; Cell Signaling Technologies, 9217 or 9275), anti-total AKT (rabbit polyclonal 1:1000; Cell Signaling Technologies, 9272S), anti-phospho-ERK1/2 (rabbit polyclonal 1:1000; Cell Signaling Technologies, 9101S), anti-total ERK1/2 (rabbit polyclonal 1:1000; Cell Signaling Technologies, 9102S). Membranes were washed three times with TBS-T for 10 min and incubated with horseradish peroxidase conjugated secondary anti-mouse IgG (1:2000, Cell Signaling Technologies, 7076) or anti-rabbit IgG (1:2000, Cell Signaling Technologies, 7074) in 5% nonfat dry milk/TBS-T for 1 h at room temperature followed by three more wash cycles with TBS-T. Blots were visualized using a ChemiDoc Imaging System (Bio-Rad, 12003153) after applying chemiluminescent substrate (Thermo Scientific, 34578, 34096) Densitometry was performed using Fiji software 2.14.0/1.54f through inversion of chemiluminescence image. All protein bands were confirmed to be at the expected molecular weight by using a molecular weight ladder (Bio-Rad, 1610394).

### Nanoparticle Tracking Analysis

The size and concentration of isolated EVs were analyzed using a NanoSight NS300 (Malvern Panalytical) as previously described (Almanza *et al*., 2018, Sadri *et al*., 2022). Briefly, EVs were diluted in sterile-filtered DPBS to yield 30-100 particles per frame, and the camera level was adjusted accordingly with the detection threshold set at 10. Samples were then injected through a low volume flow cell at a constant flow rate at room temperature. Each sample was measured in triplicate. Malvern NTA analytical software version 3.4 was used for capture and analysis.

The ZetaView QUATT (Particle Metrix) was used as a second method to determine the concentration and the diameter of EVs. Samples were diluted until approximately 500 particles per frame were observed using light scatter detection. In PT mice, fluorescing snEVs were measured at 25°C using a laser wavelength of 488 nm, 500 nm long pass light filter, shutter speed of 100, and sensitivity set to 98. Total EVs were measured at 25°C using a laser with a wavelength of 488 nm light filter, shutter speed of 150, and sensitivity set to 80 (Rupert, *et al*., 2017).

### Transmission electron microscopy

Copper mesh grids (400; Ted Pella, Inc., 01754-F) were subjected to a glow discharge using Pelco EasiGlow (Ted Pella, Inc.) and placed in a 10 μL droplet of EV-containing solution, with 5 × 10^10^ particles/mL, for 5 min. Grids were washed and stained with 2% uranyl acetate (Ladd Research, 23620). Excess uranyl acetate was removed by carefully dabbing the grid against blotting paper (Tisch Scientific, 1001-085). The grids were then left to dry overnight. Images were captured with a JEM-1400Plus (JEOL Ltd.) microscope operating at 80K eV using a Gatan OneView 4kx4k bottom mounted camera.

### Cryogenic electron microscopy

The Thermo Scientific Vitrobot Mark IV was used for sample plunge freezing. A house-made holey carbon TEM grid was first glow discharged (25 mM, 30 sec) using PELCO easiGlow™ Glow Discharge Cleaning System. Then, a 4 µL sample was applied on the grid, plunged into liquid ethane, and transferred to liquid nitrogen with a 5 second blotting time and +10 force (Marr *et al*. 2014).

The plunge-frozen TEM grid was loaded into a Thermo Scientific Glacios electron microscope (200kV) and imaged with a Gatan K3 direct electron detector. Serial EM was used to control the microscope and the detector in low dose mode at either 11,000x or 45,000x nominal magnification.

### CIPPN mouse model

PTX (5 mg; Sigma, T7191) was diluted in a 1:1 solvent with 200 proof ethanol (Decon Labs Inc., 2716) and Cremophor (EMD Millipore Corp., 238470) to make a 6 mg/ml stock solution, which was further diluted 1:15 in 8 mM sodium chloride saline (Singla *et. al.* 2002). PTX was administered by intraperitoneal injection. 4 mg/kg of PTX was injected every other day for a total of 5 doses, 20 mg/kg cumulative dose. Painful peripheral neuropathy was determined based on pain-related behaviors (Toma *et. al.* 2022).

To test mechanical hypersensitivity, mice were placed in a plexiglass box (22×22×14 cm) on a metal grid floor and acclimated to the behavior testing facility for at least 60 minutes. Mechanical sensitivity was tested by applying 0.04 to 4 g Von Frey Filaments (Stoelting, 58011) to the plantar surface of the left hind paw. Filaments were presented consecutively ascending or descending using the up-down method previously described and modified for the mouse (Chaplan *et al*., 1994, Poplawski *et al*., 2018). The filament that caused paw withdrawal 50% of the time was determined as the paw withdrawal threshold (50% PWT). All experiments were performed by an investigator blinded to the mouse identity.

To test cold hypersensitivity, experiments were performed as previously described (Toma *et al*., 2017) with modifications. Mice were placed in a plexiglass box (22 × 22 × 14 cm) on a metal grid floor and acclimated for at least 30 min before testing. 20 μL of acetone was projected onto the plantar surface of the hind paw from a pipette tip. Time spent licking, shaking, and holding the hind paw was recorded by a stopwatch for 2 min.

### DIA LC-MS/MS Proteomics

Samples were reconstituted in 200 μL of 6 M guanidine-HCl and boiled for 10 min followed by 5 min of cooling at room temperature. The boiling and cooling cycle was repeated a total of 3 times. The proteins were precipitated with addition of methanol to final volume of 90 µl followed by vortexing and centrifugation at maximum speed on a benchtop microfuge (14,000 rpm) for 10 min. The soluble fraction was removed by flipping the tube onto an absorbent surface and tapping to remove any liquid. The pellet was suspended in 200 μL of 8 M urea made in 100 mM Tris pH 8.0. TCEP and chloro-acetamide were added to final concentrations of 10 mM and 40 mM, respectively, and vortexed for 5 min. 3 volumes of 50 mM Tris pH 8.0 were added to the sample to reduce the final urea concentration to 2 M. Trypsin was added and incubated at 37°C for 12 h. The solution was then acidified using TFA (0.5% TFA final concentration) and mixed. The sample was desalted using C18-StageTips (Thermo Fisher) as described by the manufacturer protocol. The peptide concentration of sample was measured using BCA. 0.20 μg of each sample were used for DIA analysis.

To perfoirm DIA mass spectrometry, 1 μg of peptides for each sample (BCA quantification) was analyzed by ultra-high pressure liquid chromatography (UPLC) coupled with tandem mass spectrometry (LC-MS/MS) using nano-spray ionization. The nanospray ionization experiments were performed using a TimsTOF HT pro hybrid mass spectrometer (Bruker) interfaced with nano-scale reversed-phase UPLC (EVOSEP ONE). Evosep method 30 samples per day was utilized using a 15 cm × 150 μm reverse-phase column packed with 1.5 μm C18-beads (PepSep, Bruker) at 58°C. The analytical columns were connected with a fused silica ID emitter (10 μm ID; Bruker Daltonics) inside a nanoelectrospray ion source (Captive spray source; Bruker). The mobile phases comprised 0.1% FA as solution A and 0.1% FA/99.9% ACN as solution B. Settings for the TimsTOF Pro HT followed PASEF method for standard proteomics. The values for mobility-dependent collision energy ramping were set to 95 eV at an inversed reduced mobility (1/*k*_0_) of 1.6 V s/cm^2^ and 23 eV at 0.73 V s/cm^2^. Collision energies were linearly interpolated between these two 1/*k*_0_ values and kept constant above or below. No merging of TIMS scans was performed. Target intensity per individual DIA-PASEF precursor was set to 20,000. The scan range was set between 0.6 and 1.6 V s/cm^2^ with a ramp time of 166 ms. 10 PASEF MS/MS scans were triggered per cycle (1.17 s) with a maximum of seven precursors per mobilogram. Precursor ions in an *m*/*z* range between 100 and 1700 with charge states ≥3+ and ≤8+ were selected for fragmentation. Active exclusion was enabled for 0.4 minutes (mass width 0.015 Th, 1/*k*_0_ width 0.015 V s/cm^2^). Peptide DIA quantifications were carried out using Peaks Studio 12 (Bioinformatics solutions Inc.).

### Mouse dorsal root ganglia primary cultures

Primary dorsal root ganglion (DRG) neuron cultures were established from adult C57BL/6J mice, following previously published protocols (Wang *et al*., 2022) and modified for mice (Sleigh *et al*., 2019). Briefly, DRGs were digested with collagenase A (Sigma Aldrich, 10103578001) and glial cells were disassociated by 0.125% trypsin (Gibco, 15140122). Cells were cultured in Neurobasal medium (Thermo Fisher, A3582901) supplemented with 100 U/mL penicillin and 100 μg/mL streptomycin, 2% heat-inactivated FBS, 2% heat-inactivated HS, 2% B27 (Thermo Fisher, 17504044), 2 mM GlutaMAX (Thermo Fisher, 35050061), and 100 ng/mL β-nerve growth factor (β-NGF; Sigma Aldrich, N2513). Cells were pelleted at 800 x g for 5 min at room temperature. Cells were cultured onto 6-well plates pre-coated with 10 µg/mL poly-D-lysine (PDL) and 10 µg/mL laminin. After one day, the culture medium was replaced, supplemented with 10 µM cytosine β-D-arabinofuranoside (AraC; Sigma Aldrich, C6645) overnight to inhibit glial cell proliferation, after which the culture medium was replaced once again. Thereafter, the medium was exchanged every other day and supplemented with fresh β-NGF.

### Cell Signaling in primary sensory neurons

DRG neuron-enriched cultures were transferred to serum-free medium for 45 min prior to treatment with snEVs (1×10^9^ particles/well) isolated from either healthy or neuropathic mice for 5 or 30 min. Cells were rinsed with ice-cold DPBS and extracted in RIPA lysis buffer by scraping plate. Extracts were centrifuged at 15,000 x g for 10 min at 4°C, and the supernatant was collected. Equal amounts of protein from cell extracts (20 µg), as determined by BCA, were subjected to immunoblotting as described above.

### Fluorescence-based quantification of tGFP in nerve tissue extracts and snEVs

Tissue samples were homogenized in radio-immunoprecipitation assay buffer (RIPA; Sigma Aldrich, R0278) supplemented with 1% protease inhibitor (Thermo Fisher, 78429) and 1% phosphatase inhibitor (Thermo Fisher, 78420) using a Sonic Dismembrator Model 500 (Thermo Fisher). Protein concentration was quantified by microBCA Protein Assay Kit (Thermo Fisher, 23235) and equivalent amounts of protein were loaded into the wells of a 96 well plate (Thermo Fisher, 12-566-70). A standard curve was obtained using known concentrations of fluorescein (Sigma Aldrich, F6377) in 10 mM Tris buffer, pH 10. The relative fluorescent values (RFV) were measured using a Spectramax-iD3 (excitation/emission wavelengths at 462/502 nm) and analyzed with Softmax Pro v.7 software. Background was determined by using solution only-controls.

### Immunofluorescence (IF) microscopy

Sciatic nerves from PT mice were isolated, fixed with 4% paraformaldehyde (PFA) and fibers were teased apart after removal of the epineurium or transferred to cryostat embedding medium (O.C.T.; Tissue Plus, 4568) and placed over dry ice until solidified. Frozen nerves were sectioned by cryostat to 20 μm thick sections. Nerve tissues were fixed onto glass micro slides (VWR, 48311-703), followed by permeabilization using 0.4% Triton-X100 (Fisher Bioreagents, 9002-93-1) in 1% horse serum (HS, Sigma Aldrich, H1270). Slides were blocked in 5% HS diluted in DPBS for 30 min at room temperature. Immunofluorescent staining was performed with anti-tGFP VHH (ChromoTek, tbt) conjugated with Mix-n-Stain™ CF-568 antibody labeling kit (Sigma Aldrich, MX568S20) for 2 h at room temperature or CellMask (Thermo Fisher, C10046) diluted in UltraPure water (Invitrogen, 10-977-023) for 10 min. Mounting medium containing 4’,6-diamadino-2-phenylindole (DAPI; Abcam, ab104139) was used.

### Primary mouse SC cultures

Primary mouse SCs were isolated from adult mice as previously described (Martellucci *et al*., 2025, Poplawski *et al*., 2018, Azzouz *et al*., 1996). Briefly, mice were first subjected to sciatic nerve crush injury. After 7 days, nerve tissue at the crush site and distal to the injury was collected. The epineurium was removed and digested with collagenase D1 (Sigma Aldrich, SCR139). Nerves were titurated gently 10-15 times, as described above (see Isolation of sciatic nerve-derived EVs). Isolated cells were cultured in DMEM, supplemented with 10% heat-inactivated FBS, 100 U/mL penicillin and 100 μg/mL streptomycin (Gibco, 15140122), 1 nM neuregulin-1 (NRG1, R&D Systems, 9875-NR-050), and 2 µM forskolin (Cell Signaling Technologies, 3828S), in 60 mm wells pre-coated with 100 µg/mL PDL (Sigma Aldrich, P6407) and 10 µg/mL laminin (Gibco, 23017015). When cellular confluence was approximately 85%, the cultures were treated first with anti-Thy1.2 antibody (Thermo Fisher, 14-0903-82) and subsequently with rabbit complement (Sigma Aldrich, S7764). Cells were passaged no more than 3 times before use in experiments. There were no significant metabolic or morphological changes in passage numbers 1-3.

### SC IF microscopy

Cultured primary SCs were cultured on Millicell EZ slides (Millipore Sigma, PEZGS0416), fixed in 4% PFA, permeabilized using 0.1% Triton-X100, and blocked with 5% BSA. Fixed cells were incubated with anti-SOX10 (Thermo Fisher, MA5-32335) diluted 1:500 in blocking solution over night at 4°C. The secondary anti-rabbit Alexa Fluor 594 conjugated antibody (Invitrogen, A21442) was then added for 45 min at room temperature. Mounting medium containing DAPI was used for mounting the cover slip (Avantor, 48393-106). Images were taken using an Olympus IX73 microscope using CellSens Dimension 3.2 software (Olympus).

#### Super resolution microscopy with direct stochastic optical reconstruction (dSTORM)

snEVs were characterized using the Nanoimager S from Oxford Nanoimaging Inc. (ONI), as previously described using the propriety reagents by ONI’s Application Kit^TM^: EV Profiler 2 (900-00079) (Senesi *et al*., 2024; Gebara *et al*., 2022). Samples were normalized so that an equivalent number of EVs were used per acquisition (∼6.9 × 10^10^ particles/mL). EVs were captured using either biotinylated phosphatidylserine antibody or biotinylated hCD9 antibody. Both Pan-EV and human reactive CD9 detection antibody were applied in conjunction to each lane of the assay chip and incubated for 70 minutes.

dSTORM imaging buffer A was first brought to room temperature. 100 uL of a 1:100 working reagent was prepared using one part imaging buffer B (chilled) to 99 parts imaging buffer A. Contents were carefully mixed with a pipet to ensure adequate mixture prior to application to all lanes on assay chip. Assay chips were imaged using the Nanoimager S with 640 nm laser was set to 50% for 1,000 frames and 488 nm laser was set to 100% for 3,000 frames. All images were analyzed using the AutoEV workflow acquisition provided on ONI’s Collaborative Discovery (CODI) online analysis platform (www.alto.codi.bio).

### Statistical Analysis

Statistical analysis was performed using GraphPad Prism 10.2.0 (GraphPad Software). All results are presented as the mean ± S.E.M. Comparisons between two groups were performed using two-tailed unpaired T-tests. In some cases, non-parametric data was analyzed by a Mann Whitney U test. When comparing more than two groups, a one-way or two-way ANOVA and Tukey’s *post hoc* test was performed. P<0.05 was considered statistically significant. (* p < .05; ** p < .01; *** p < .001; **** p < .0001).

## Results

### Isolation of EVs from mouse sciatic nerves

In order to isolate peripheral nerve EVs with minimal contaminating cell fragments, adult mouse sciatic nerves (SN) were extracted with a collagenase and dispase blend (C1D), a method previously shown to leave cells intact (Martellucci *et al*., 2024). RIPA extracts of SN were isolated and compared as a control. Nerve extracts were analyzed by immunoblotting to detect cellular β-actin, a biomarker of cell fragments. β-actin was detected only in the RIPA extracts and not in the C1D extracts, indicating that C1D digests are largely free of cellular components (**Fig. 1A**).

**Figure 1.**
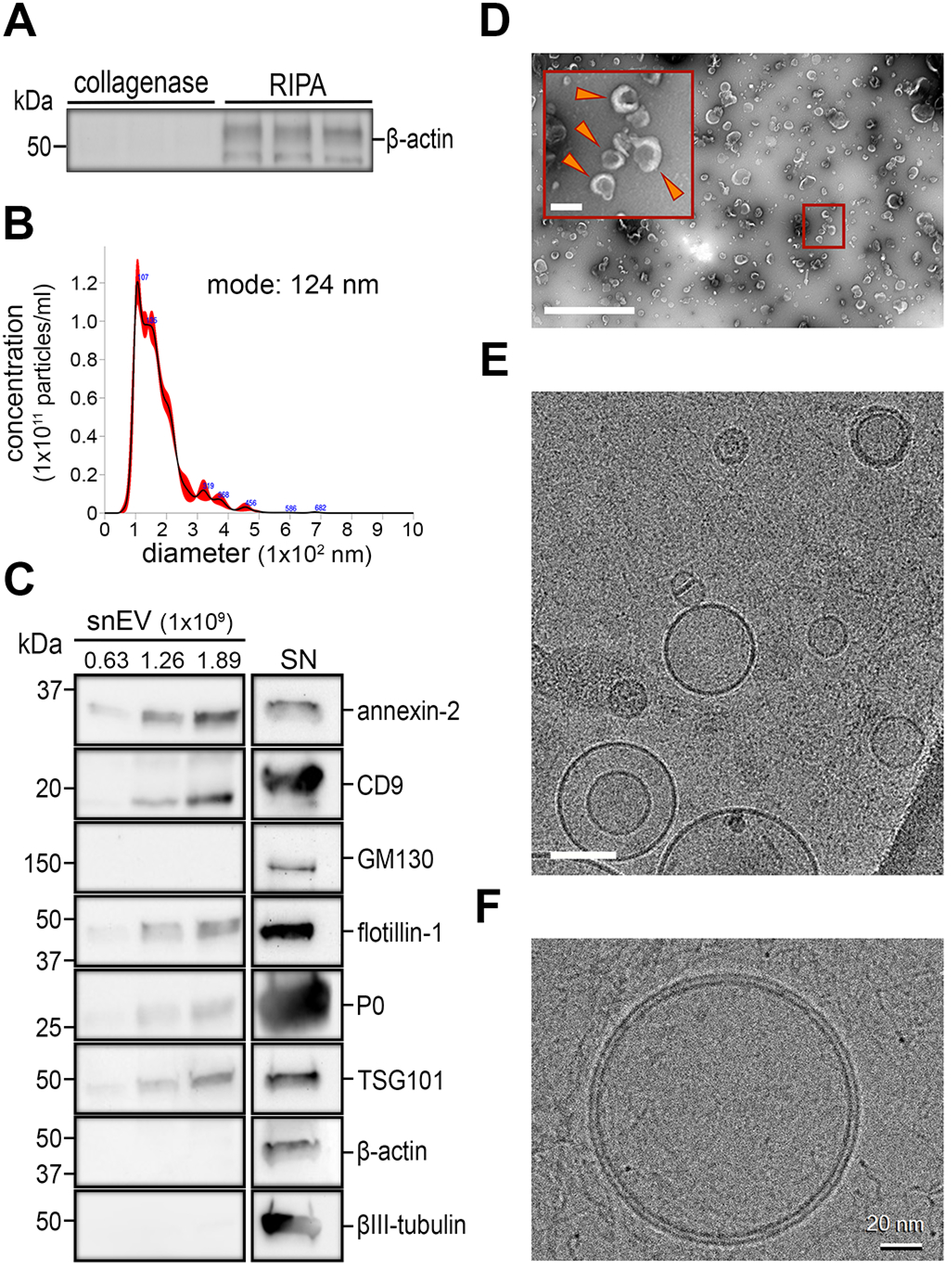
Characterization of extracellular vesicles (EVs) isolated from sciatic nerves (SN) in mice. **A.** Immunoblot analysis of collagenase (CID) or RIPA digested extracts of SN tissue to detect cellular protein β-actin. n=3/group. **B.** Representative nanoparticle tracking analysis (NTA) histogram of snEVs with reported modal diameter (124 nm) and concentration (1.4 ×10^11^ particles/mL), acquired by NanoSight NS300. n=3 technical replicate captures, 60 second recordings each. **C.** Representative immunoblots of snEVs with dose dependent concentrations of snEVs, acquired by NTA (left), whole sciatic nerve lysates (10 µg) (right). EV biomarkers detected: annexin-A2, CD9, flotillin, and TSG101. GM130 is a Golgi marker, the neuron specific cytoskeleton protein βIII-tubulin and cellular protein ß-actin were used as negative controls. The SC biomarker, myelin protein zero (P0) was found to be present in snEV samples. **D.** Representative transmission electron microscopy (TEM) image of negatively stained snEVs. Arrow heads highlight individual snEVs displaying cup-like morphology. Scale bar, 200 nm (inset), 2 µm (bottom). **E.** Representative cryogenic electron microscopy (cryo-EM) image showing the heterogenous population of snEVs. Optical zoom, 11,000x; Scale bar, 100 nm. **F.** Representative cryo-EM image of a single snEV showcasing the molecular architecture of a lipid bilayer. Optical zoom, 45,000 x; Scale bar, 20 nm.

Sciatic nerve EVs were isolated from C1D extracts by centrifugation and SEC (**Supplemental Fig. 1**). Nanoparticle tracking analysis (NTA) demonstrated snEVs were present in the C1D extracts and successfully harvested from an intact multicellular tissue (**Fig. 1B**). The snEVs were heterogeneous and distributed into a single major peak with a modal diameter of ∼124 ± 17 nm. To test whether the harvested snEVs included known EV biomarkers, immunoblot analysis was performed. Increasing concentrations of snEV particles (0.63, 1.26 and 1.89 × 10^9^ particles, corresponding to 1.04, 2.08 and 3.12 µg of snEV protein, respectively) were loaded into each well and compared to mouse SN tissue (10 µg total protein) harvested in RIPA buffer. Proteins detected in the snEV preparation, which are known to be associated with EVs, included: annexin-A2, the tetraspanin protein and glia biomarker, CD9, the lipid raft-associated protein, flotillin, and the ESCRT cytoplasmic marker, TSG101 (**Fig. 1C**). GM130, a Golgi apparatus marker that is reported to be excluded from EVs (Merianda *et al*., 2009; Lötvall *et al*., 2014), was absent from the snEV preparation. The cellular cytoskeletal proteins, β-actin and neuron specific βIII-tubulin were also not detected in snEVs. The SC biomarker, P0, was present in snEVs, demonstrating that at least a subpopulation of the snEVs was derived from SCs. EVs are produced by SCs in culture (Sadri *et al*., 2022); however, EV production by SCs *in vivo* has not been previously demonstrated.

Transmission electron microscopy was performed to examine the snEVs isolated using C1D and purified by SEC. The snEVs demonstrated the expected cup-shaped morphology observed by sample preparation for TEM (**Fig. 1D**). Since this does not represent their native structure and particle identity (Kurtjak *et al*., 2022), we performed cryo-EM to examine the structure of hydrated snEVs. Images of snEVs from healthy sciatic nerves revealed variation in size and structural complexity (multi-layer) (**Fig. 1E**). The snEVs had intact lipid bilayers, which is characteristic of EVs (**Fig. 1F**). These results confirm that the C1D-based EV isolation procedure did not damage snEVs.

### Development of the CIPPN mouse model and the impact on size or concentration of snEVs

To determine whether snEVs are altered by CIPPN, we utilized a previously described model system (Li *et al*., 2015) with some modifications. Baseline behavioral testing for mechanical hypersensitivity and cold allodynia was performed in 4-month-old C57BL/6 mice. The mice were then treated with PTX or vehicle control every other day for ten days. Pain-related behaviors were measured weekly beginning immediately after cessation of PTX treatment, for a total of 4 weeks. A sustained decrease in PWTs, an indicator of mechanical hypersensitivity, was observed throughout the time course after PTX treatment (**Supplement Fig. 2A**), replicating previous results (Li *et al*., 2015). Decreased PWTs report increased mechanical hypersensitivity induced by PTX. PTX treatment also induced significant cold hypersensitivity immediately after cessation of PTX administration that worsened after two weeks (**Supplement Fig. 2B**). Vehicle controls manifested pain related behaviors similar to naïve mice (Poplawski *et al*., 2018), thus, naïve mice were considered healthy mice and used through our studies.

Cryo-EM images of healthy snEVs (snEV_H_) and snEVs from PTX-treated (snEV_NP_) mice revealed similar native, hydrated structure morphology of EVs with intact lipid bilayers and some noticeable heterogeneities in size and internal cargo (**Fig. 2A, B**). Single nanoparticle tracking analysis (Zetaview) and NTA (NanoSight) of snEVs isolated from healthy and PTX-treated (neuropathic) nerves immediately after completion of PTX administration (week 0) demonstrated that the concentration (**Fig. 2C, D**) and size (**Fig. 2E-G**) of snEVs was similar before and after PTX treatment.

**Figure 2.**
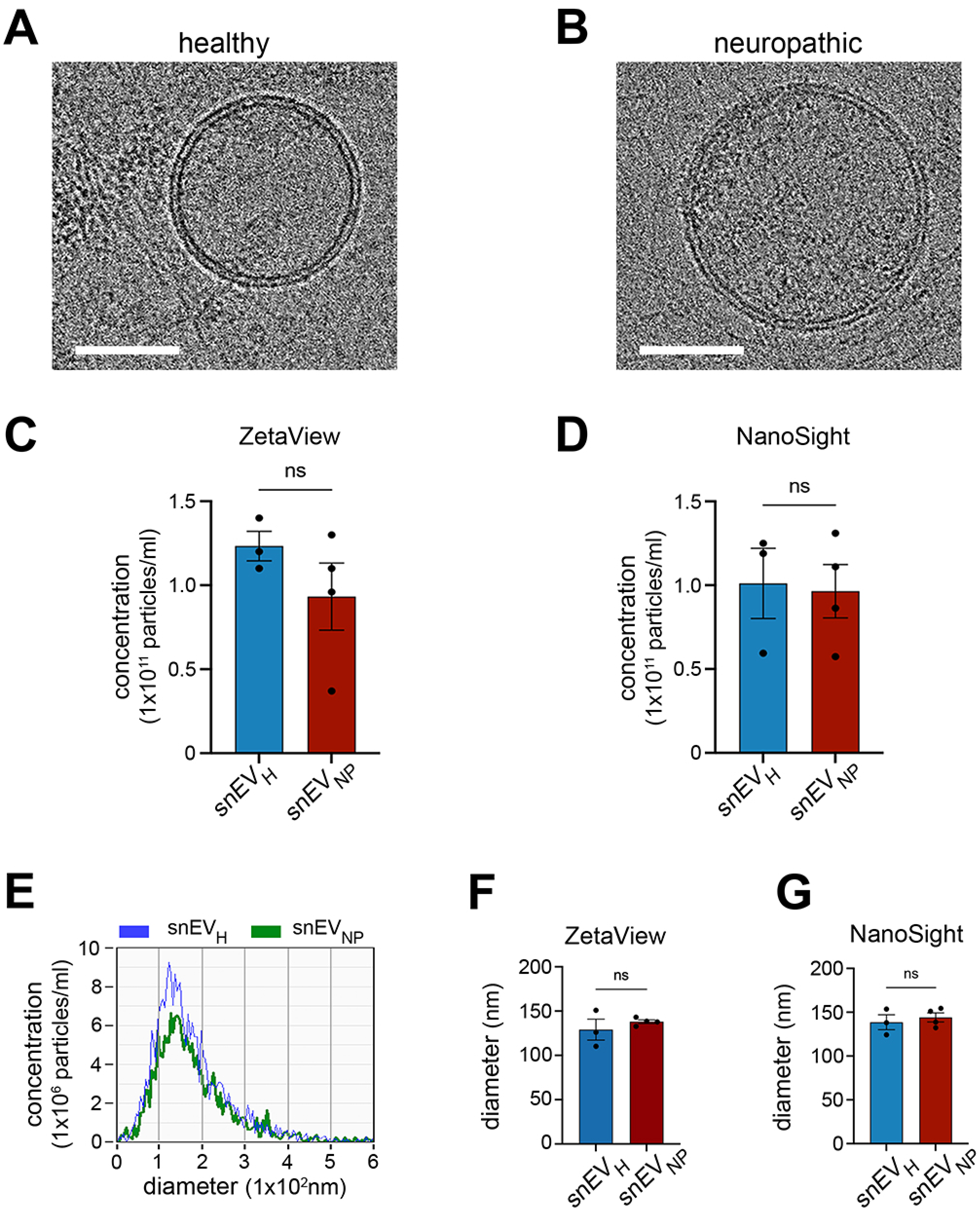
Paclitaxel induces a painful peripheral neuropathy that does not alter overall snEV concentration or size. **A.** Representative cryo-EM image highlighting the structure of a single snEV isolated from healthy mice (snEVH). Optical zoom, 45,000x; Scale bar, 50 nm. **B.** Representative cryo-EM image highlighting the structure of a single snEV isolated from neuropathic mice (snEVNP) mice. Optical zoom, 45,000 x; Scale bar, 50 nm. **C-D**. Average concentration of snEVH and snEVNP determined by NTA acquired by (C) ZetaView QUATT and (D) NanoSite NS300, N=3-4 independent replications. Data are expressed as mean ± SEM; ns, not significant. **E.** Representative histograms acquired by single particle tracking analyses (ZetaView Quatt) of snEVH (blue) and snEVNP (green), acquired by ZetaView QUATT, displaying similar size distribution. **F-G.** Modal diameter of snEVH and snEVNP determined and acquired by (F) ZetaView QUATT, N=4 independent replications and (G) NanoSight NS300, N=3-4 independent replications. Data are expressed as mean ± SEM; ns, not significant.

### snEVs isolated from mice with CIPPN have a unique and altered proteome

To determine whether snEVs isolated from CIPPN mice have alterations in protein cargo, we conducted a highly sensitive quantitative proteomic analysis of healthy and neuropathic snEVs in biological quadruplicates. Principal component analysis showed separation of healthy and neuropathic samples with greater sample variation in healthy snEVs, suggesting EV hetero-geneity. DIA-based LC-MS/MS identified more than 2000 proteins in each snEV phenotype. Differential protein abundance analysis of neuropathic versus healthy samples showed enrichment of 47 proteins and depletion of 70 proteins in neuropathic snEVs. Protein identification and quantification were performed using a PEAKS workflow on DIA LC-MS/MS data, and proteins were filtered using −10logP confidence scores and a global false discovery rate threshold (**Fig. 3A; Supplement Table 1,2**). Annexin-A2, a validated extracellular vesicle biomarker (Grieve *et al*., 2012), was amongst the most abundant proteins detected in both healthy and neuropathic snEVs, and was validated by immunoblotting (**Supplement Fig. 3A**), further confirming EV enrichment. Extracellular vesicles biomarkers, such as flotillin, were also present at similar levels. These findings are consistent with our single nanoparticle tracking analysis showing no difference in nanoparticle concentration between the snEV groups.

**Figure 3.**
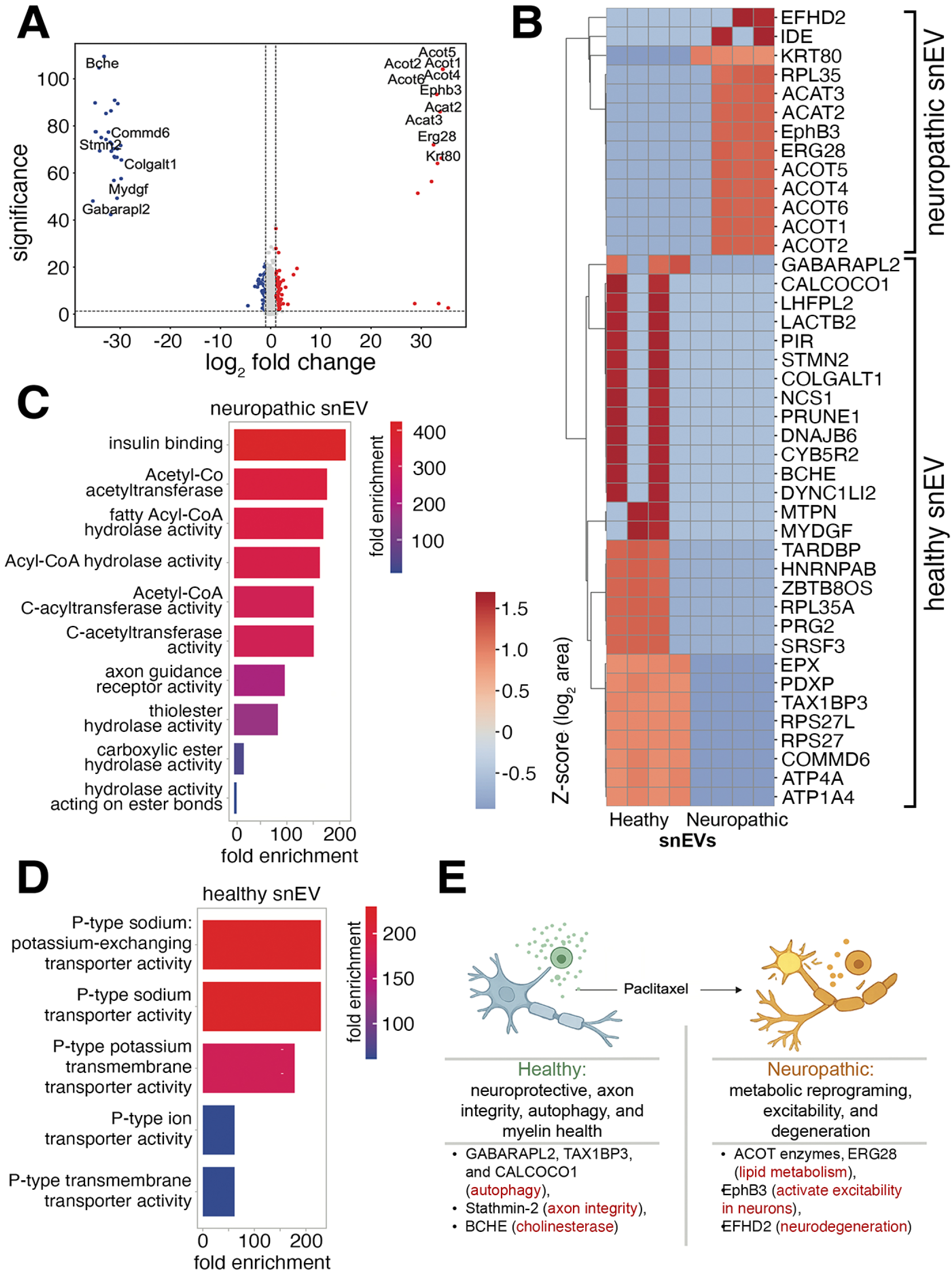
Unique and differential level proteins of snEVs from neuropathic versus healthy mice. **A.** Volcano plot showing unique and differentially expressed proteins between snEVs isolated from healthy (snEVH) and neuropathic (snEVNP) mice. Proteins with log₂ fold change ≥ 1 and significance ≥ 15 are highlighted in red (up in neuropathic) or blue (up in healthy). **B.** Heatmap (Z-score of log₂-transformed areas) displaying expression patterns of uniquely expressed proteins across snEVH and snEVNP samples. Unsupervised clustering shows groups of proteins based on expression patterns in the two experimental groups. **C-D.** Bar plots show GO Molecular Function enrichment analysis of proteins unique to snEVH (C) or snEVNP (D). **E.** Schematic summary highlighting biological implications: snEVH carry proteins supporting neuroprotection (autophagy, axon integrity, myelin stability), whereas snEVNP are enriched in proteins associated with neuropathic pain mechanisms, including metabolic reprogramming, neuronal excitability, and degeneration.

Importantly, 29 proteins were identified as unique to healthy snEVs, and 13 proteins were unique to neuropathic snEVs (**Fig. 3B**). Gene ontology (GO) analyses of proteins uniquely detected in healthy snEVs revealed enrichment of proteins related to axon integrity and proteins that are essential for maintaining resting membrane potential and transport of cations, crucial for cellular homeostasis. In contrast, neuropathic snEVs exhibited significant enrichment of proteins related to fatty acid metabolism and neuronal cell signaling when analyzed by GO Molecular function (**Fig. 3C, D**). We noted that healthy snEVs uniquely expressed ATG8-family proteins related to autophagy including GABARAPL2, TAX1BP3 and CALCOCO2. Previous work on EV heterogeneity has shown an integration of autophagy and EV protein composition (Xu *et al*., 2018).

Healthy snEVs also expressed abundant proteins associated with peripheral nerve health and axon integrity including, stathmin-2 (STMN2), cholinesterase (BCHE) and procollagen galactosyltransferase (COLGALT1). We also observed proteins involved in maintaining cell survival (MYDGF) and integrity and health of myelin in mature nerves (COMMD6). By contrast, neuropathic snEVs contained several acyl-CoA thioesterases (ACOTs) that are essential for controlling the levels of fatty acyl-CoAs to maintain metabolic health. They also expressed cell surface receptors and ligands (EphB3, FGF1, AGPAT1, NRADD) known to activate excitability in sensory neurons, and or play a role in neurodegeneration (EFHD2) (**Supplemental Table 1; Supplemental Fig. 3B**). Several Rab proteins (RAB1A and RAB14) known to be involved in EV biogenesis were downregulated in neuropathic EVs (**Supplemental Table 2; Supplemental Fig. 3B**). A summary of snEV phenotype is shown in **Fig. 3E** and indicates specific proteins in healthy snEV proteins which contribute to neuroprotection and processes important for homeostasis (autophagy) whereas neuropathic snEVs contain protein cargo that is associated with metabolic reprogramming and neuronal excitability.

### snEVs activate cell-signaling in primary sensory neurons

We established primary cultures of sensory neurons from mouse DRGs to test whether snEVs isolated from SNs activate cell-signaling. Our previous studies demonstrated that EVs isolated from human plasma directly induce activation of ERK1/2 in PC12 cells (Gonias *et al*., 2022). Wild type primary sensory neuron cultures were highly enriched in βIII-tubulin, but not the glia marker, SOX10 (**Fig. 4A**). Sensory neurons were treated with snEVs isolated from healthy mice or neuropathic mice for 5 or 30 min. Healthy snEVs did not activate ERK1/2 after 5 min (determined as ratio of phosphorylated ERK1/2 (pERK1/2) to total ERK1/2 (tERK1/2)), whereas neuropathic snEVs activated ERK1/2 at 5 min (**Fig. 4B).**

**Figure 4.**
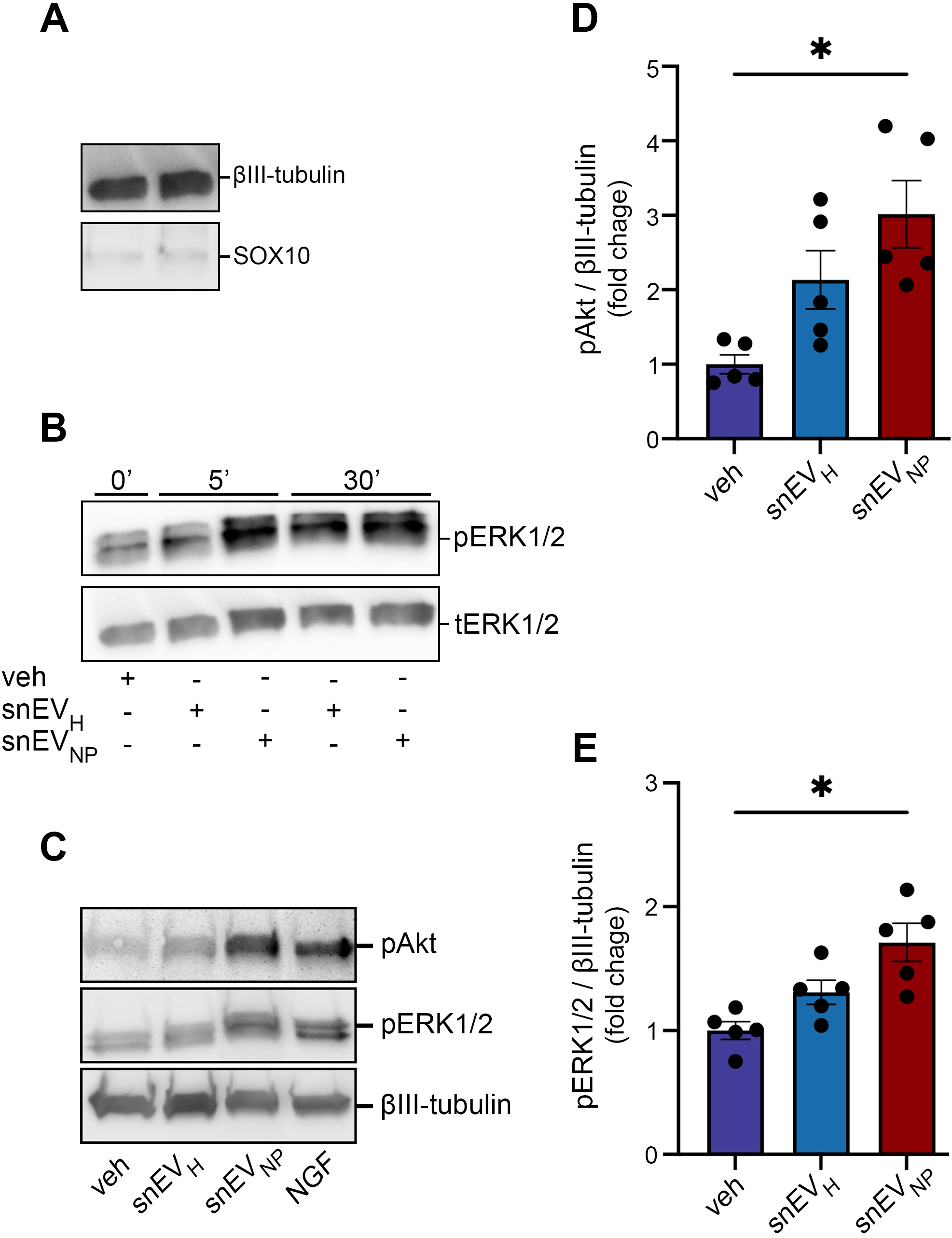
Neuropathic snEVs acutely activate pain associated cell signaling pathways in primary cultures sensory neurons. **A.** Representative immunoblots show enrichment of βIII-tubulin and SOX10 in primary sensory neuron cultures sourced from dorsal root ganglion (DRG) of wild type mice. **B.** Representative immunoblots show an increase in phospho-ERK1/2 (pERK1/2) and total ERK1/2 (tERK1/2) in primary mouse DRG neurons at 5 and 30 minutes of incubation with SEC elution buffer (veh), snEVH or snEVNP. Cultures were treated with 1 × 10^9^ particles/well or vehicle, all at an equivalent volume. **C.** Representative immunoblot analyses of pAKT, pERK1/2, and βIII-tubulin in DRG derived primary sensory nerve culture lysates; 20 μg/lane. Culture lysates were incubated with SEC elution buffer (veh), snEVH, snEVNP, or NGF for 5 minutes per treatment, all at an equivalent volume. N=5 independently prepared primary cultures. **D-E.** Densitometry quantification of immunoblots of (D) pAKT and (E) pERK1/2 as a ratio to βIII-tubulin and normalized to vehicle. Data is expressed as the mean ± SEM; N=5 independently prepared primary cultures. Data was analyzed using Kruskal-Wallis test with Dunn’s multiple comparison test. * *P*<0.05.

Next, we compared the cell-signaling responses (activation of AKT and ERK1/2) elicited by snEVs isolated from healthy and neuropathic mice. snEVs were added to primary sensory neuron cultures for 5 min. Neuropathic, but not healthy snEVs, activated AKT within 5 min (**Fig. 4C**). This study also replicates the results regarding ERK1/2 activation shown in **Fig. 4B**. Densitometric analyses of snEV signaling revealed a greater than 3-fold activation of AKT (**Fig. 4D**) and greater than 1.5-fold activation of ERK1/2 (**Fig. 4E**) by neuropathic snEVs.

### Identification of SC snEVs isolated from the sciatic nerve

To specifically examine SC EV subpopulations in snEVs, we generated P0-Cre-TIGER (PT) transgenic mice. These mice express endogenous tGFP fused to hCD9 selectively in SCs. To validate this new mouse model system, we examined tGFP expression in various extracts of organs by fluorometry (**Fig. 5A**). Sciatic nerves demonstrated abundant fluorescence, as anticipated. By contrast, brain, lung, liver, and spinal cord showed no detectable fluorescence.

**Figure 5.**
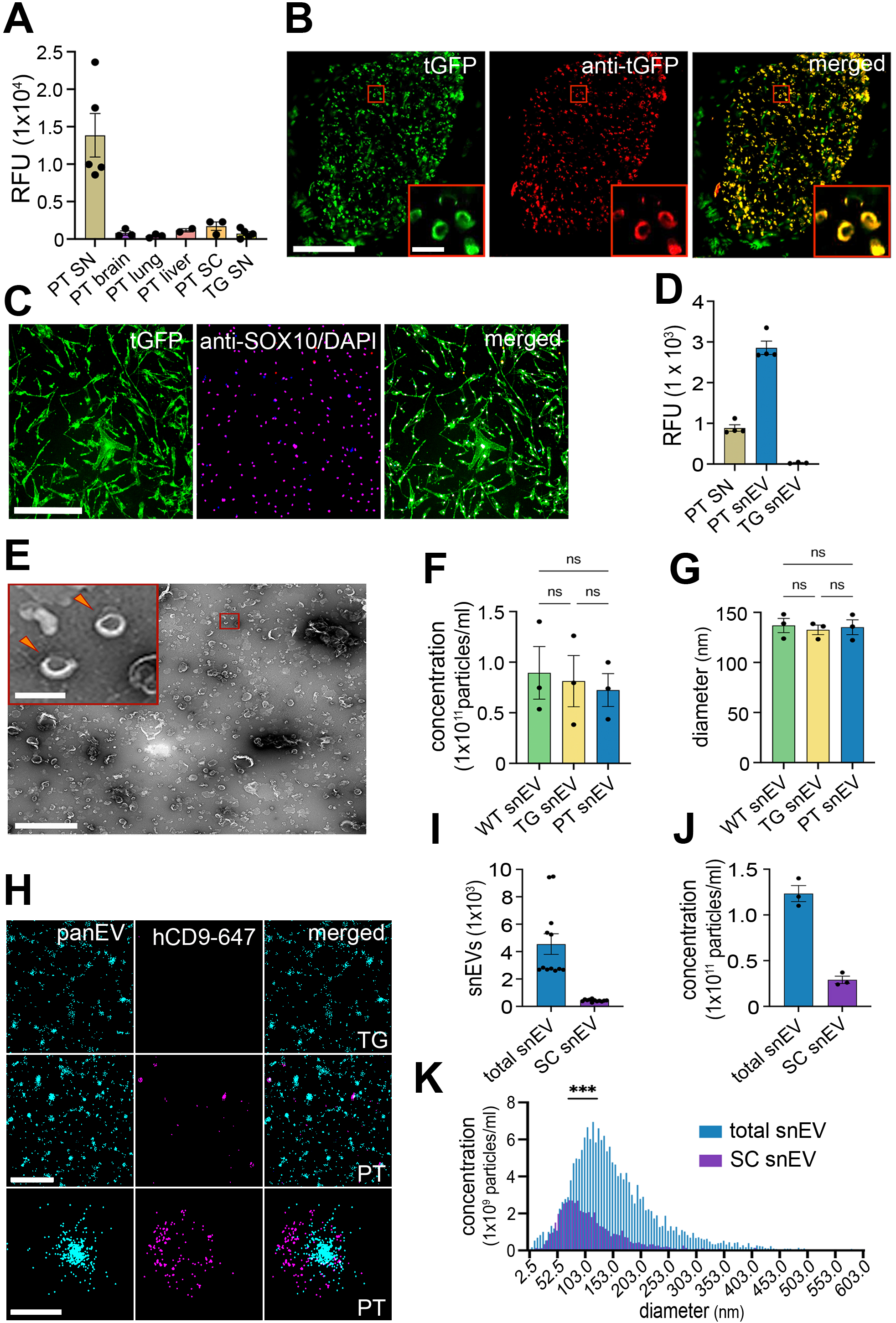
SC from P0-*Cre* x TIGER (PT) mice produce EVs expressing a tGFP-hCD9 construct. **A.** Quantitative analyses of relative fluorescence (RFU) in tissues from P0-Cre-TIGER (PT) mice. Fluorescence of sciatic nerve (SN), brain, lung, liver spinal cord (SC) and Cre-littermate control sciatic nerve (TG SN) was measured. Data are expressed as mean (RFU/µg protein) ± SEM. N=2-5 mice/group. **B.** Fluorescent and immunofluorescent (IF) images of transverse section of PT sciatic nerve (SN) showing endogenous expression of tGFP (green), fluorescent-labeled tGFP identified by an anti-tGFP nanobody (red), and merged images (yellow). Scale bar, 100 µm (left) and 10 μm (inset). **C.** Fluorescent and IF images of a primary SC culture isolated from PT mice expressing endogenous tGFP (green), fluorescent labeled SOX10 (red) and DAPI (blue), and merged images. Scale bar, 500 µm. **D.** Quantification of relative fluorescence of SN lysate isolated from PT mice, snEVs isolated from PT mice, and snEVs isolated from TG mice. All samples were normalized to equal protein concentration prior to fluorescent measurement acquisitions. Data are expressed as mean ± SEM. N=4 independent biological replicates/group. **E.** Representative transmission electron microscopy (TEM) image of negatively stained snEVs isolated from PT mice. Arrow heads highlight individual snEVs displaying cup-like morphology typical of EVs Scale bar, 200 nm (inset), 2 µm (bottom). **F-G.** Concentration (F) and modal diameter (G) of snEVs isolated from wild type (WT), TG, and PT mice acquired by NTA using NanoSight NS300. Data is expressed as the mean ± SEM, N=3 independent snEV preparations/group. Data are analyzed by a one-way ANOVA and Tukey’s *post hoc* test (ns, not significant). **H.** Representative images of total snEVs and SC snEVs isolated by phosphatidylserine (PS) capture using super-high resolution direct stochastic optical reconstruction microscopy (dSTORM). snEVs were stained using lipophilic dye, PanEV (cyan) and human specific CD9 (hCD9-647) (purple). Top row: wide field of view of snEV isolated from TG mice. Middle row: wide field of view of snEV isolated from PT mice. Scale bar, 1 µm. Bottom row: super-high-resolution image of a single double-positive (PanEV + hCD9-647) snEV isolated from PT mice. Scale bar, 200 nm. **I.** Quantification of single positive (PanEV only) and double-positive (PanEV + hCD9-647) snEVs isolated from PT mice using phosphatidylserine capture method and analyzed by CODI software. Data is expressed as the mean ± SEM. n=12 technical replicates of N=2 independent experiments or replicates. **J.** Concentration of total snEVs (light scatter) and tGFP positive snEVs (fluorescent), acquired by ZetaView QUATT nanoparticle analyses. Data is expressed as the mean ± SEM. N=3 independent snEV preparations. **K.** Overlayed size distribution histograms of total snEV and tGFP positive snEVs, acquired by single nanoparticle fluorescence analyses. Results are expressed as the mean. N=4 independent snEV preparations. Asterick indicates significant differences in modal diameter of SC snEVs when compared to the total snEV population, analyzed using an unpaired T-test test (*** *P*<0.001).

Next, we examined transverse sciatic nerve sections by fluoresence spectroscopy and IF microscopy. Endogenous tGFP was clearly imaged by fluorescence microscopy in the absence of antibodies (**Fig. 5B).** The signal was located in multiple crescent-shaped SCs (green) as anticipated. Using an anti-tGFP nanobody (red), co-localization of tGFP antigen with the endogenous tGFP fluorescence signal was observed, demonstrating the known morphology of SCs. The tGFP was localized in the SC cytoplasm of myelinated fibers, but not in the cell bodies or Satellite cells in the DRG (**Supplement Fig. 4A**). Preparations of teased fibers revealed that tGFP was (localized to paranodal junctions in sciatic nerves (**Supplement Fig. 4B, C**), consistent with the location of EV release in myelinated fibers (Samara *et al*., 2013).

To confirm that tGFP was selectively expressed by SCs in PT mice, SCs were isolated from SNs of adult animals and established in primary culture. SCs were fixed and immunolabled with SOX10 (a SC biomarker) and DAPI. Sox10 and tGFP co-localized to >95% of DAPI stained cells, confirming their SC identity. (**Fig. 5C**).

Next, we prepared CID extracts from snEVs from PT mice and control Cre negative littermate mice (TG mice). Using fluorimetry, PT mouse snEVs extracts show highly enriched fluoresence in tGFP, compared with the equivalent protein load of TG mouse snEV extracts, as anticipated. The fluorescence signal generated by PT mouse snEVs was 3-fold greater than extracts of intact PT SN (**Fig. 5D**). These results confirm that at least a fraction of the snEVs isolated from healthy peripheral nerve are derived from SCs.

Transmission electron microscopy was performed to examine PT mouse snEVs. Similar to wild-type mouse snEVs (**Fig. 1D**), PT mouse snEVs were modestly heterogeneous in size and frequently demonstrated cup-shaped morphology (**Fig. 5E**). The yield of snEVs from PT and TG mice was similar to wild type snEVs (approximately 7.5 × 10^10^ snEVs/mL) (**Fig. 5F**) and the size distribution of harvested snEVs was similar amongst the genotypes (modal diameter of approximately 136 nm) (**Fig. 5G; Supplement Fig. 5A-D**).

To characterize SC snEVs in PT mice, we initially utilized dSTORM super high-resolution microscopy. We took advantage of the fact that the outer membranes of many EVs are rich in phosphatidylserine (PS) (Gonias *et al*., 2022). snEVs from PT mice were purified by SEC and then subjected to PS immunoaffinity capture using a PS-specific antibody. PS-captured snEVs were subsequently labeled with a proprietary lipophilic dSTORM-compatible fluorophore Pan-EV (cyane), that identifies all EVs, and an anti-hCD9 antibody conjugated to a complementary dSTORM-compatible fluorophore (purple) (**Fig. 5H**). Importantly, the CD9 antibody only reacts with human CD9, but not mouse CD9. A representative dSTORM image of an individual snEV is shown in the bottom set of panels in **Fig. 5H**. Each individual cluster was assumed to represent a single snEV. Amongst the 4,000 clusters identified, approximately 500 were immunopositive for hCD9 and thus identified as SC snEVs. By this method, SC snEVs represented 10-15% of the total snEV preparation (**Fig. 5I**).

As a second approach for identifying and quantifying SC snEVs from sciatic nerves of PT mice, we utilized single fluorescent nanoparticle analysis. With this method, total snEVs are detected by light scatter and endogenously fluorescent, tGFP-expressing snEVs are detected separately. In this analysis, the total number of snEVs isolated from 10 nerves was 1.2 ± 0.1 × 10^11^ per mL (**Fig. 5J**). The corresponding number of tGFP-expressing snEVs was 2.9 ± 0.4 × 10^10^ per mL. Thus, compared with dSTORM, single fluorescent nanoparticle analysis revealed the fraction of SC snEVs was slightly higher (24%), which is not surprising given its sensitivity range.

When we analyzed the size of EVs in the total versus fluorescent population, the SC snEVs were smaller than the overall snEV population (**Fig. 5K**). The modal diameter of SC snEVs was approximately 75 nm, suggesting that these snEVs may be enriched in exosomes, which tend to be smaller (30-150 nm) than other EVs (Greening and Simpson, 2018).

### PTX regulates SC snEV subpopulations in painful peripheral neuropathy

Although we saw no changes in the overall concentration of snEVs isolated from healthy versus CIPPN mice (**Fig. 2C,D**), we hypothesized that the population of SC derived snEVs may change. To test this hypothesis, we subjected snEVs from healthy and neuropathic PT mice to dSTORM. We measured hCD9-expressing snEVs after PS immunoaffinity capture. Representative dSTORM images showed a decrease in the number of snEVs observed following PTX treatment (**Fig. 6A, left panels**), suggesting that the abundance of SC-derived snEVs was decreased. To confirm that the labeling antibody was targeting human CD9, we examined snEVs isolated from Cre negative littermate control mice (TG); no SC-derived snEVs were detected (**Supplemental Fig. 6A**). At higher magnification, an individual SC snEV isolated from PTX-treated mice demonstrated increased diameter compared to controls (**Fig. 6A, right panels**). We further confirmed these findings by immunocapturing SC snEVs directly using a hCD9-specific antibody (**Supplemental Fig. 6B, C**). Overall, the results obtained using dSTORM (**Fig. 6B**) demonstrated that the fraction of SC-derived snEVs isolated from CIPPN mice was decreased by approximately 60% (**Fig. 6C**).

**Figure 6.**
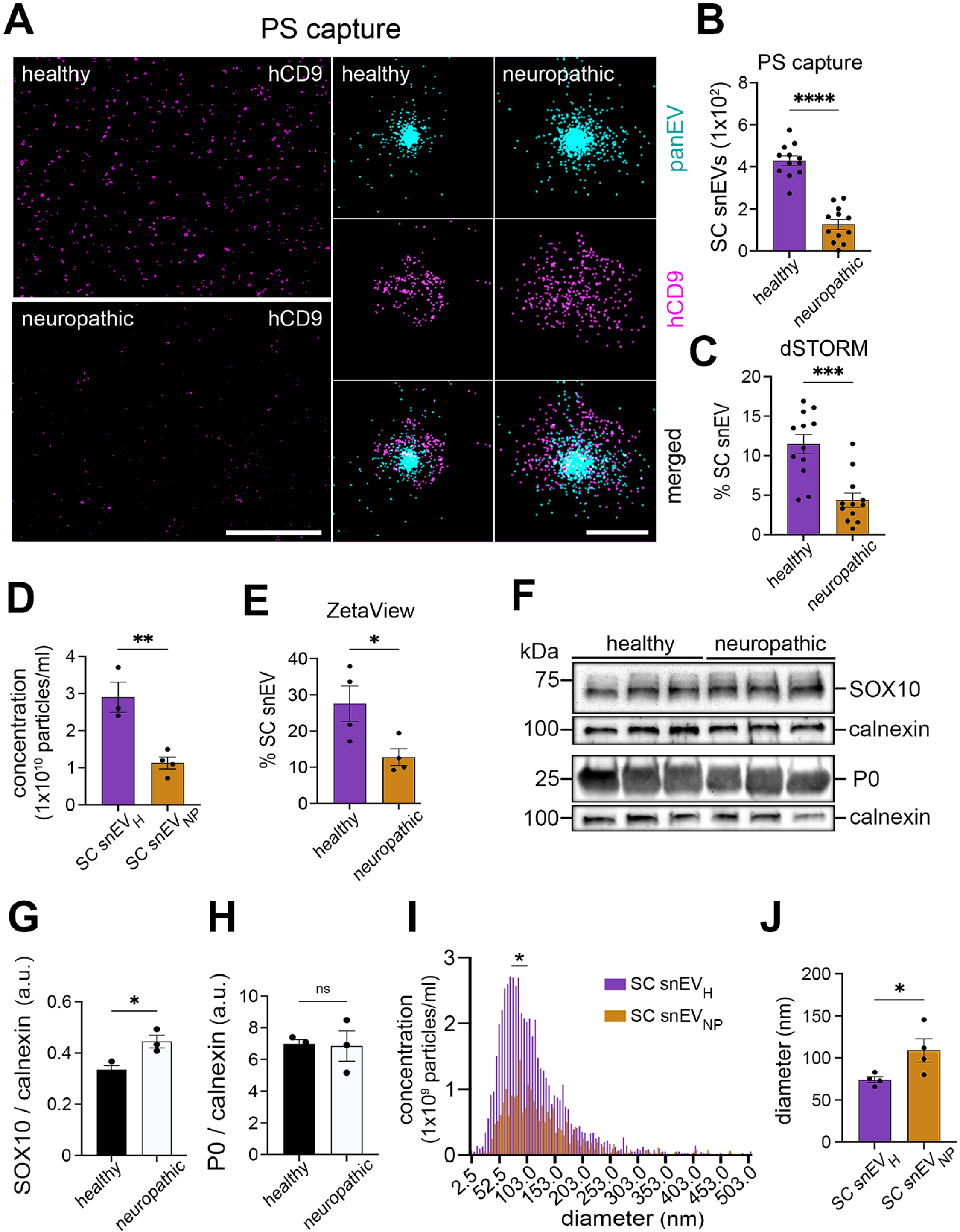
SC snEVs isolated from neuropathic mice (snEVNP) are less abundant and larger in diameter than SC snEVs from healthy mice (snEVH). **A.** (Left) representative wide field view of super-high resolution dSTORM images of SC snEVH and SC snEVNP after anti-PS immunoaffinity capture and anti-hCD9 labeling (purple); scale bar, 5 µm. (Right) representative high resolution dSTORM images of a single snEV, labeled with lipophilic dye PanEV (top; cyan), hCD9-647 (middle; purple), and merged images (bottom). Scale bar, 200 nm. **B.** Analysis of PanEV labeled snEVs that are immunopositive for hCD9-647 (SC) in healthy and neuropathic snEVs after PS capture. snEVs measuring greater than 300 nm in diameter were excluded from analysis. n=12 technical replicates of N=2 independent replications. Data are expressed as mean ± SEM; **** P<0.0001. **C.** Percentage of SC snEVs in healthy or neuropathic mice represented in total snEV populations. SC snEVs are identified by PS capture, PanEV labeling and hCD9-647 immunopositivity. n=12 technical replicates of N=2 independent replications. Data are expressed as mean ± SEM; *** P<0.001. **D.** Concentration of SC snEVH and SC snEVNP, acquired by ZetaView QUATT. N=3-4 independent replications. Data are expressed as mean ± SEM; ** P<0.01. SC snEVNP points shared with Fig. 3J. **E.** Percentage of SC snEVs in health and neuropathic mice in total snEV populations acquired by ZetaView QUATT. N=4 independent replications. Data is expressed as the mean ± SEM; * P<0.05. **F.** Immunoblots of sciatic nerve lysates from healthy and neuropathic mice immunopositive for the Schwann cell biomarkers SOX10, myelin protein zero (P0), the loading control calnexin; 10 µg protein per lane. N=3 independently prepared tissue lysates. **G**-**H**. Densitometry analysis quantifying relative band intensity of (G) SOX10 or (H) P0 relative to calnexin in panel (F). Data are expressed as the mean ± SEM. * *P*<0.05. **I.** Overlayed size distribution histograms of SC snEVH and SC snEVNP, acquired by single nanoparticle tracking analysis acquired with ZetaView QUATT. N=4 independently prepared snEV sample. Asterick indicates significant increase in modal diameter between snEVH and snEVNP; * P<0.05. **J.** Modal diameter of SC snEV_H_ and SC snEV_NP_, determined by single nanoparticle tracking analysis acquired with ZetaView QUATT. Data are expressed as mean ± SEM; N=4 independently prepared snEV samples; * *P*<0.05.

Next, we analyzed snEVs isolated from healthy or neuropathic PT mice by single fluorescent nanoparticle tracking analysis. Total EV concentration as measured by light scatter was similar in both groups (**Fig. 2C,D**), however, fluorescing nanoparticles were significantly lower in the neuropathic snEV preparations compared to healthy controls (**Fig. 6D**). This represented about a 60% decrease in SC snEVs (**Fig. 6E**), that was consistent with the dSTORM findings.

To demonstrate that PTX did not affect SC viability, we measured the SC biomarkers, SOX10 and P0, in nerves from healthy and PTX-treated mice by immunoblot analysis (**Fig. 6F**). Densitometric analyses demonstrated a slight increase in SOX10 expression after PTX treatment (**Fig. 6G**; p<0.05). Whether this is due to SC proliferation or an increase in SOX10 protein expression by existing SCs is not clear. However, P0 protein was unchanged by this dose of PTX treatment (**Fig. 6H**), suggesting there was no change in SC viability.

In addition to changes in SC snEV populations after PTX treatment, single nanoparticle fluorescent tracking analysis demonstrated a significant increase in the modal diameter of SC snEVs isolated from PTX-treated mice compared with healthy snEVs. The results of four independent studies are summarized in **Fig. 6I**. The SC snEV modal diameter significantly increased from about 70 nm in healthy mice to approximately 120 ± 25 nm in neuropathic mice (**Fig. 6J**).

## Discussion

This study highlights two technological advances; for the first time that EVs may be isolated from the interstitial spaces of healthy and neuropathic peripheral nerves and that SC-derived populations can be identified in snEVs utilizing a novel, sensitive tGFP-hCD9 reporter mouse. In addition, we also report that although snEVs are heterogenous in nature, the proteome of healthy and neuropathic snEVs, possess unique and distinct protein signatures in the two phenotypes. The proteomic landscape revealed that healthy snEVs support a neuroprotective role in glia-neuron communication, whereas snEVs isolated from chemotherapy-treated mice exhibit a neuropathic phenotype. Specifically, healthy snEVs contained numerous autophagy-associated proteins, such as GABARAPL2, TAX1BP1, and CALCOCO1, which are expressed in SCs (Gomez-Sanchez *et al*., 2015; Jang *et al*., 2015) and are essential for maintaining myelin and preserving intracellular organelle integrity. In myelinating glia, autophagy-dependent turnover regulates the selective breakdown and recycling of myelin proteins and lipids to sustain myelin function and normal neural activity (Belgrad *et al*., 2020). Additionally, autophagy activation can drive the biogenesis of specialized EVs termed “amphisomes” (Xu *et al*., 2018), which share autophagic molecular machinery (Ju *et al*., 2025).

Sciatic nerve-derived EVs from mice with CIPPN contained an abundance of ACOT enzymes. These enzymes hydrolyze fatty acyl-CoAs; in their absence, cells become more susceptible to metabolic stress, hyperexcitability, and neurotoxicity (Ellis *et al*., 2013). Given the close proximity of SCs to sensory neurons, ACOT deficient SCs are well poised to influence neighboring neurons, resulting in peripheral sensitization. snEVs from CIPPN mice uniquely expressed EphB3, an ephrin receptor found on EVs isolated from neuropathic mice following paclitaxel treatment. The Eph/ephrin family is the largest receptor tyrosine kinase family and comprises two subfamilies, A and B (Kania and Klein, 2016). This receptor-ligand dynamic allows for bidirectional signaling, meaning that both the receptor and ligand undergo phosphorylation upon binding, thereby activating downstream signaling pathways (Kania and Klein, 2016). Recently, it was demonstrated that human cancer cell-derived EVs carry EphB2 and facilitate reverse cell signaling to drive endothelial tube formation (Sato *et al*, 2019). Furthermore, ephrin B1/EphB1 and ephrin B2/EphB2 are implicated in the modulation and development of neuropathic pain (Battaglia *et al*., 2003; Kaur *et al*., 2024), as well as axon guidance in development (Parrinello *et al*., 2010). Although the involvement of EphB3 in neuropathic pain remains poorly documented, the inherent promiscuity of EPH receptors and ephrin ligands (Pasquale, 2004) underscores its potential relevance. Eph3B3 specific emergence during a pain state positions EphB3 as a prime candidate linking snEVs to the development of CIPPN. Thus, although the snEVs are heterogenous and contain snEVs from multiple cell types, they represent the tissue microenvironment, and proteomic analyses reveal a distinct and unique proteome in health and when experiencing CIPPN.

To test for functional differences between snEVs isolated from healthy and CIPPN sciatic nerve, we developed a bioassay using cultured primary DRGs enriched with sensory neurons. We compared acute cell-signaling events activated by the two snEV phenotypes. Neuropathic EVs rapidly activated ERK1/2, and AKT in primary sensory neurons. The speed of the response suggests that, at least initially, the effects of neuropathic snEVs on cell-signaling is not dependent on cargo transfer. Instead, the data support a model in which cell-signaling results from interaction of EV surface proteins with target cell receptors. We previously demonstrated a similar mechanism in which cellular prion protein in human plasma-derived EVs activated ERK1/2 in PC12 cells (Gonias *et al*., 2022). At this point, we have not definitively identified the cell surface macromolecule or macromolecules, in neuropthic snEVs, responsible for activation of cell-signaling. Furthermore, because we have not yet identified the cellular origin of neuropathic snEVs with cell-signaling activity, it remains possible that the active population is derived from other resident PNS cells or recruited inflammatory cells following PTX treatment. CIPPN induced by PTX is strongly associated with neuroinflammation (Li *et al*., 2015).

Although bulk analyses of snEVs revealed significant changes in protein cargo, the overall concentration or size of snEVs in CIPPN mice was not changed. Thus, we specifically examined a subpopulation of snEVs from SCs to determine whether these alterations were masked. Using our novel tGFP-hCD9 reporter mice, we demonstrated accumulation of tGFP in the paranodes (SC cytoplasm) of myelinated fiber architecture, which is consistent with known sites of EV release (Samara *et al*., 2013). Two robust and innovative technologies determined that 20-40% of total healthy snEVs are derived from SC snEVs. This was surprising given that >85% of the nucleated cell population in the uninjured sciatic nerve consists of SCs (Asbury, 1970; Campana *et al*., 2006). However, it is an important observation because myelination of many SCs in intact peripheral nerves adds complexity to the structure and stability of the plasma membrane, which would be expected to influence SC snEV production. It has been previously reported that glycocalyx polymers surrounding the plasma membrane of cells significantly influence EV biogenesis and release (Kuo and Paszek, 2021). We also noted that the average size of SC snEVs derived from healthy nerves were smaller when compared with the overall total healthy snEV population, which may be due to their pathway of biogenesis (Mathivanan and Simpson, 2010). EVs are classified by their size and biogenesis. For example, exosomes (30–150 nm) are smaller EVs formed within multivesicular bodies of the endosomal pathway and released upon fusion with the plasma membrane (Cocozza *et al*., 2020; Mathivanan *et al*., 2014). Based on their size, SC snEVs may be highly enriched with exosomes, and is consistent with their involvement in intercellular communication with neurons (Court and Alvarez, 2016).

Notably, PTX treatment significantly decreased SC-derived snEV concentrations without inducing detectable SC death. We hypothesize that this reduction disrupts normal glial-neuronal communication, thereby driving the development and maintenance of peripheral sensitization (Devor et al., 1992). This aligns with previous findings showing systemic and local administration of SC EVs isolated from primary rodent conditioned medium alleviates pain-related behaviors (Wang *et al*., 2020; Sadri *et al*., 2022). Because PTX-induced SC dysfunction is linked to changes in autophagy (Maiarù *et al*., 2025) and given the close relationship between autophagy and EV biogenesis (Xu *et al*., 2018), an imbalance in these pathways may cause neuropathic SCs to release fewer, larger EVs. Furthermore, our proteomic analysis revealed that neuropathic EVs contain Rab proteins, critical for vesicle trafficking. Secreting these Rab proteins into EVs likely depletes the biogenesis machinery within the parent cell (Kumar *et al*., 2024), prolonging the biogenesis period. This extended timeframe may increase the size of neuropathic SC snEVs, enhancing their cargo capacity while impairing their ability to communicate within the local nerve microenvironment. Taken together, our results suggest that neuropathic alterations in SC snEVs contribute to chronic pain states such as CIPPN. Testing this hypothesis directly will require new technologies capable of sorting SC snEV populations that remain bioactive and support significant recovery.

Collectively, we propose that a unique proteome in snEVs derived from CIPPN mice may contribute to peripheral sensitization and ongoing neuropathic pain (Devor *et al*., 1992). These proteomic results provide numerous targets for further studies to determine responsible mechanisms. Furthermore, changes in SC snEV size and concentration may underlie other structural and composition differences in snEVs that influence pain states. In future studies, the technologies described herein may be expanded to allow purification of SC-derived snEVs from *in vivo* sources such as intact peripheral nerves. Understanding the activities of EVs from specific cell types *in vivo* remains an important objective.

## Author Contributions

**Miles Vecchitto**: conceptualization, methodology, writing – original draft, data curation, formal analysis, investigation. **Gail Funk**: data curation, formal analysis, methodology. **Zixuan**: investigation, formal analysis, methodology. **Takahito Arai**: methodology, data curation, formal analysis. **Stefano Martellucci**: methodology, investigation, formal analysis, conceptualization. **Saptarshi Sinha**: methodology, investigation, formal analysis, resources. **Aaron Tran**: data curation, formal analysis, methodology. **Masaki Norimoto**: conceptualization, investigation, supervision. **Majid Ghassamian**: data curation, investigation, methodolgy. **Pradipta Ghosh**: investigation, formal analysis, resources, writing – review and editing. **Steve L. Gonias:** conceptualization, formal analysis, writing and restructuring, reviewing and editing. **Wendy Campana**: conceptualization, investigation, validation, formal analysis, visualization, funding acquisition, writing the original draft, writing, review and editing, resources, funding acquisition, project administration, supervision.

## Acknowledgements

We’d like to thank ONI for their assistance with high resolution microscopy and dSTORM studies. We would also like to thank Dr. Takuma Oligiri for his help with immunoblotting EV biomarkers and Dr. Rachel Howard-Tills for copy editing the paper. Transmission electron microcopy images were taken in the Cellular and Molecular Medicine Electron Microscopy Core (UCSD-CMM-EM Core, RRID: SCR_022039). Cryo-EM was performed at the UC Davis Biological Electron Microscopy Facility.

