## Supplemental Materials for "Peripheral nerve derived extracellular vesicles are dynamically regulated in chemotherapy induced painful neuropathy"

**Supplemental Figure 1. Size exclusion chromatography (SEC) isolates snEVs from other proteins**

A. Fractions 3-20 (30 µL/lane) isolated using Izon qEVoriginal SEC columns were subjected to SDS-PAGE and silver staining. snEVs are collected after the default buffer volume (DBV), fractions 1-6, and are shown to be enriched in the purification collection volume (PCV), fractions 7-10, consistent with the manufacturer’s data.
B. Concentration of snEVs of fractions 4-13, collected by SEC during, determined by NTA acquired using NanoSight NS300. n=3 technical replicate.
C. Immunoblot of fractions 6-11 (30 µL/lane), collected by SEC during snEV isolation. Fractions 8 and 9 of the PCVs were immunopositive for the SC biomarker P0, and EV biomarker annexin-A2 with fraction 8 having the strongest signal. Thus, fraction 8 was used throughout the remainder of the study.

**Supplemental Figure 2. Paclitaxel induces neuropathy measured by paw withdrawal threshold and cold response.**

A. Mechanical hypersensitivity in mice after a cumulative dose of PTX (20mg/kg) administered over the course of ten days by determining the 50% paw withdrawal threshold (PWT) by the up-down method using von Frey filaments. N=4 mice/group. Data are expressed as mean ± SEM. * P<0.05, ** P<0.01.
B. Cold allodynia in mice measured by recording total seconds (s) of flicking, licking, and holding of paw after stimulation with acetone. N=4 mice/group. Data are expressed as mean ± SEM. * P<0.05, ** P<0.01, *** P<0.001, **** P<0.0001.

**Supplemental Figure 3. Differentially regulated proteins of snEV_H_ and snEV_NP_.**

A. Representative immunoblot of annexin-A2 expressed in healthy (snEV_H_) and neuropathic (snEV_NP_) snEVs.
B. Volcano plot showing differentially expressed proteins between snEV_H_ and snEV_NP_ after removing unique proteins in each phenotype. Proteins with log₂ fold change ≥ 1 and significance ≥ 15 are highlighted in red (up in neuropathic) or blue (up in healthy).

**Supplemental Figure 4. Molecular architecture of sciatic nerves reveal expression of tGFP in paranode regions of healthy mice.**

A. Cross section (10 μm) of L5 DRG isolated from a PT mouse stained using CellMask 647 (red) and DAPI (blue) showcasing the absence of tGFP (green) in neuronal cell bodies and satellite cells, but present in myelinating (SC) fibers.
B. Longitudinal sections (20 μm) of teased sciatic nerve fibers isolated from a PT mouse stained using CellMask 647 (red) and DAPI (blue) highlighting the location of endogenous tGFP (green).
C. Intact myelinated fiber isolated from a sciatic nerve of a PT mouse. Arrowhead (orange) indicates the node of Ranvier. Arrows (green) highlight endogenous tGFP located in paranode of Schwann cells.

**Supplemental Figure 5. Size distribution histograms of snEVs from wild type (WT), Cre- littermate (TG), and P0-Cre-tGFP (PT) mice by NTA.**

A-C. Representative NTA size distribution histograms of snEV from (A) WT, (B) TG, and (C) PT mice, acquired by NanoSight NS300 demonstrating similarities in size distribution of snEVs amongst genotypes. N=3 replicates.

**D**. Representative overlaying size distribution histograms of snEV, light scatter (blue) and

fluorescence (green) single nanoparticle tracking analyses acquired by ZetaView QUATT

of one snEV sample isolated from PT mice.

**Supplemental Figure 6. SC derived snEVs are less abundant in mice following treatment with PTX.**

A. Quantification of PanEV labeling and anti-hCD9-647 immunopositivity of Cre- littermate (TG) snEVs after immunoaffinity capture with anti-phosphatidylserine (PS) and analyzed by CODI software. n=12 technical replicates of N=2 independent replications. Data are expressed as mean ± SEM.
B. Representative dSTORM images of SC derived snEV_H_ (top) and snEV_NP_ (bottom) after hCD9 immune capture followed by anti-CD9 conjugated with a dSTORM compatable fluorophore (purple) and analyzed by CODI software. Scale bar, 5 μm.
C. Analysis of hCD9 positive SC snEV_H_ and SC snEV_NP_ after hCD9 immunoaffinity capture. snEVs measuring greater than 300 nm diameter were excluded from analysis. n=12 technical replicates of N=2 independent replications. Data are expressed as mean ± SEM; ** P<0.01.

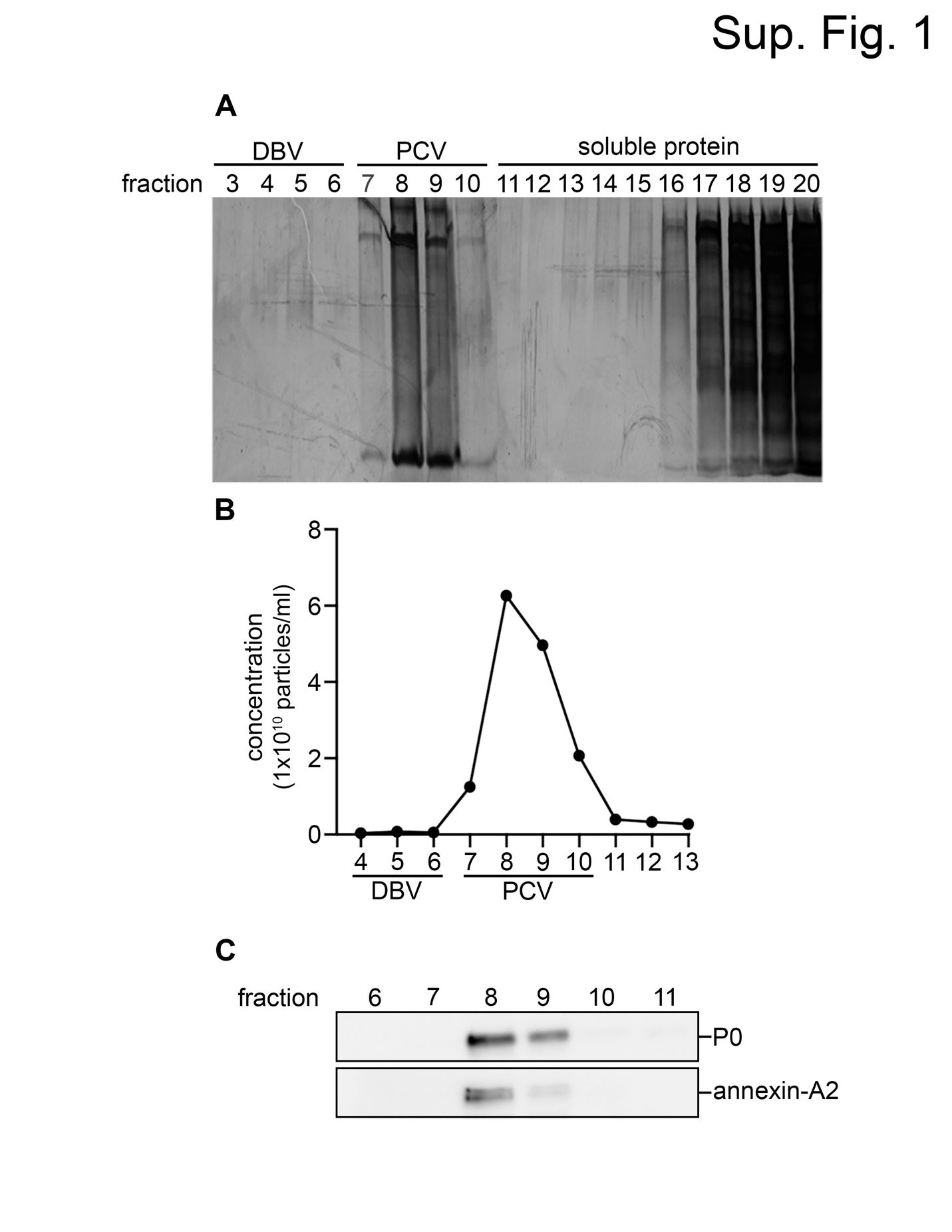

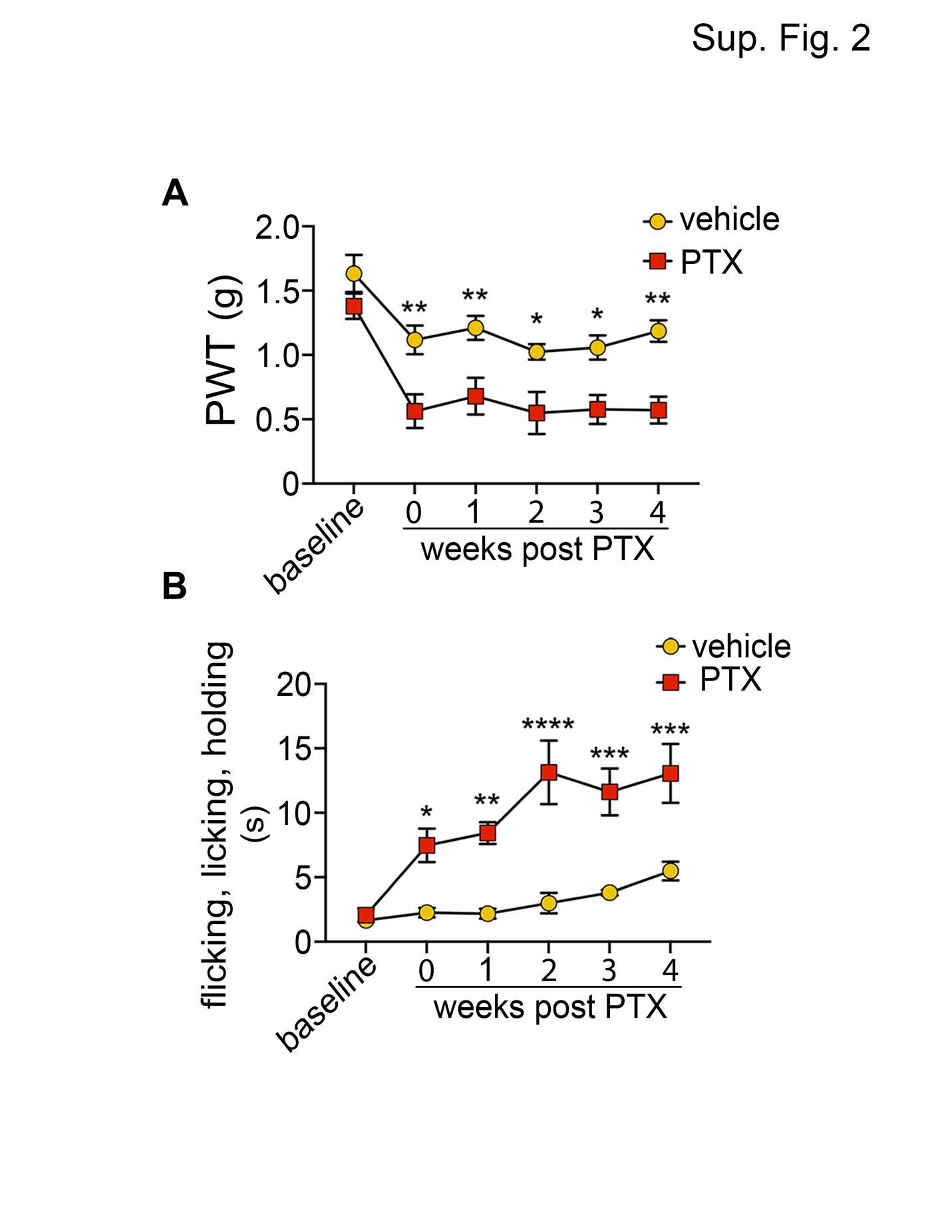

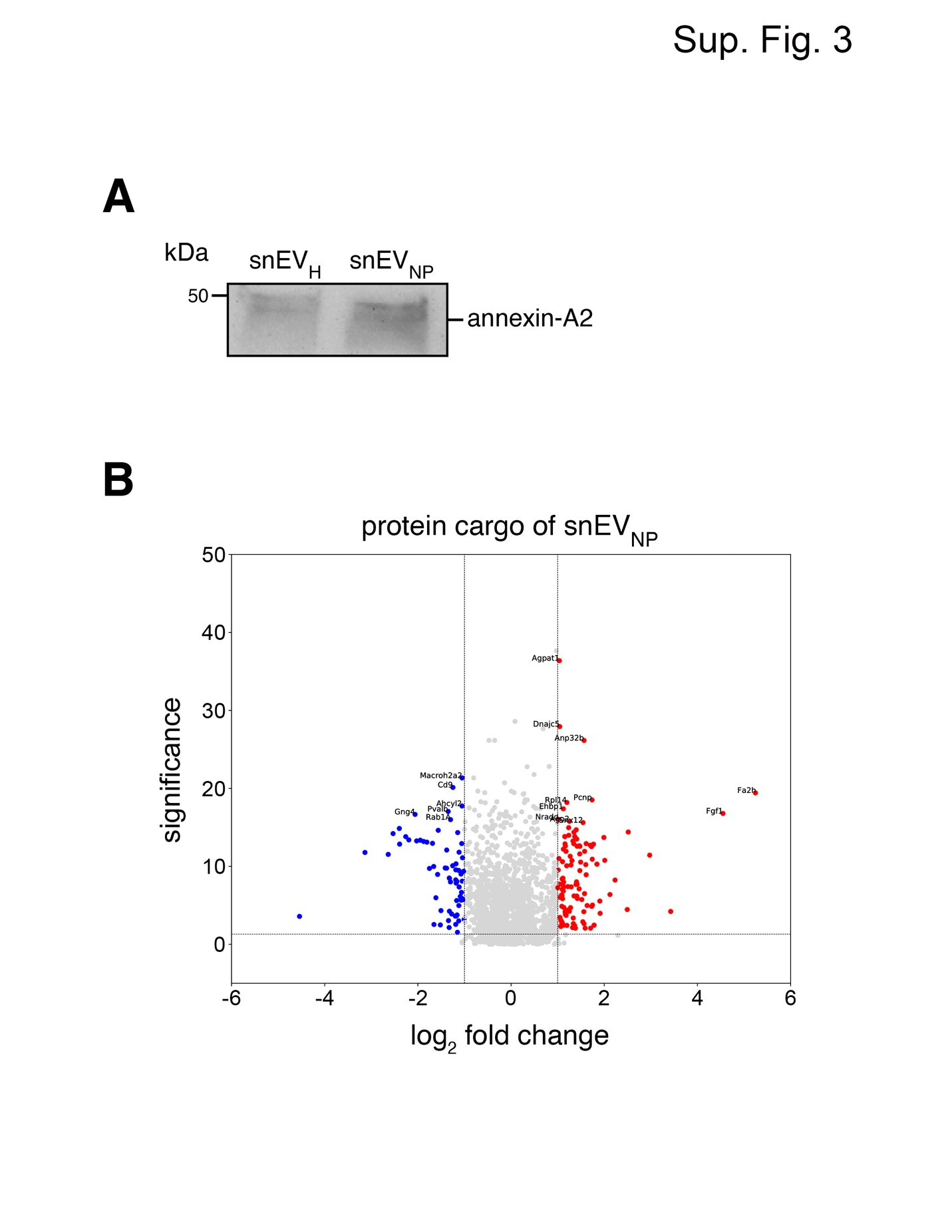

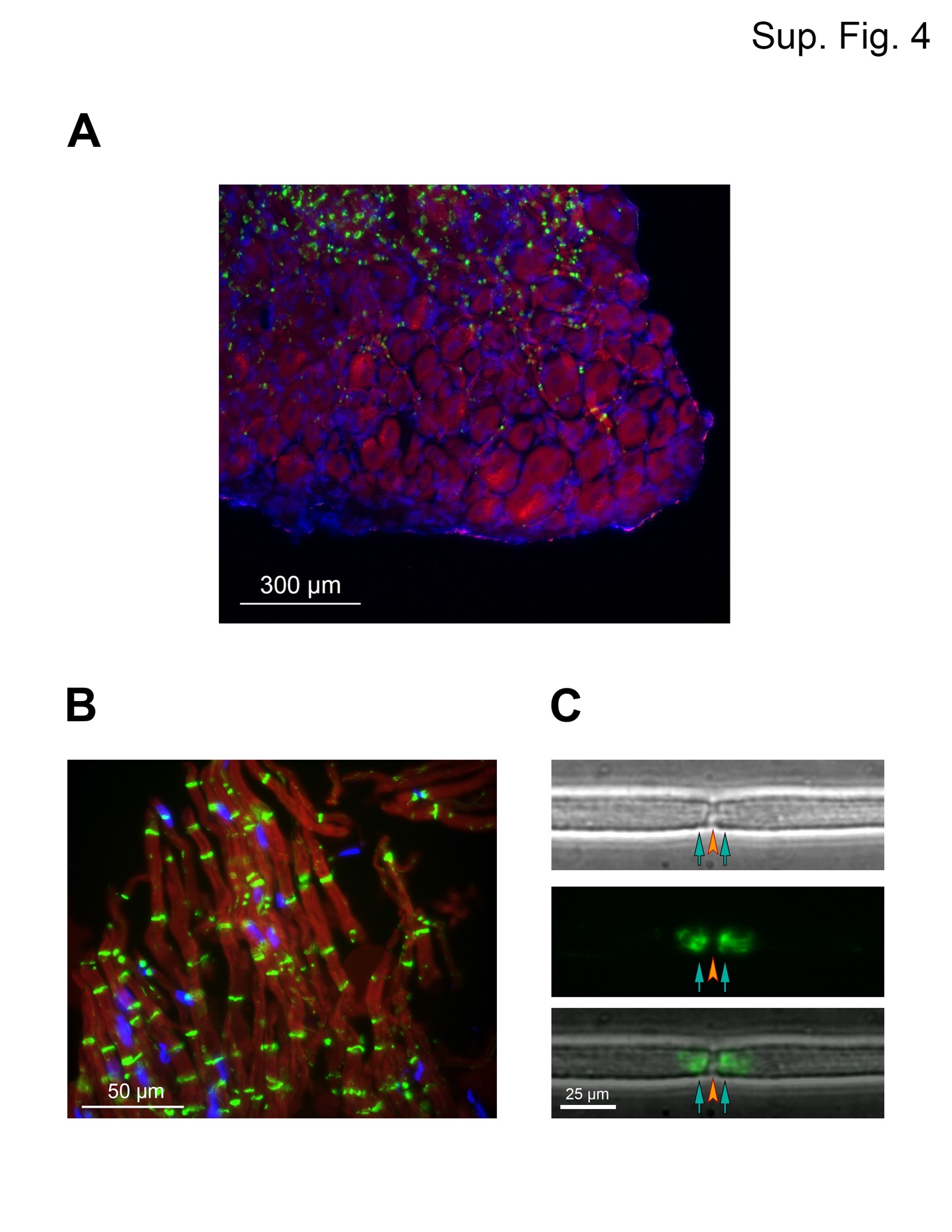

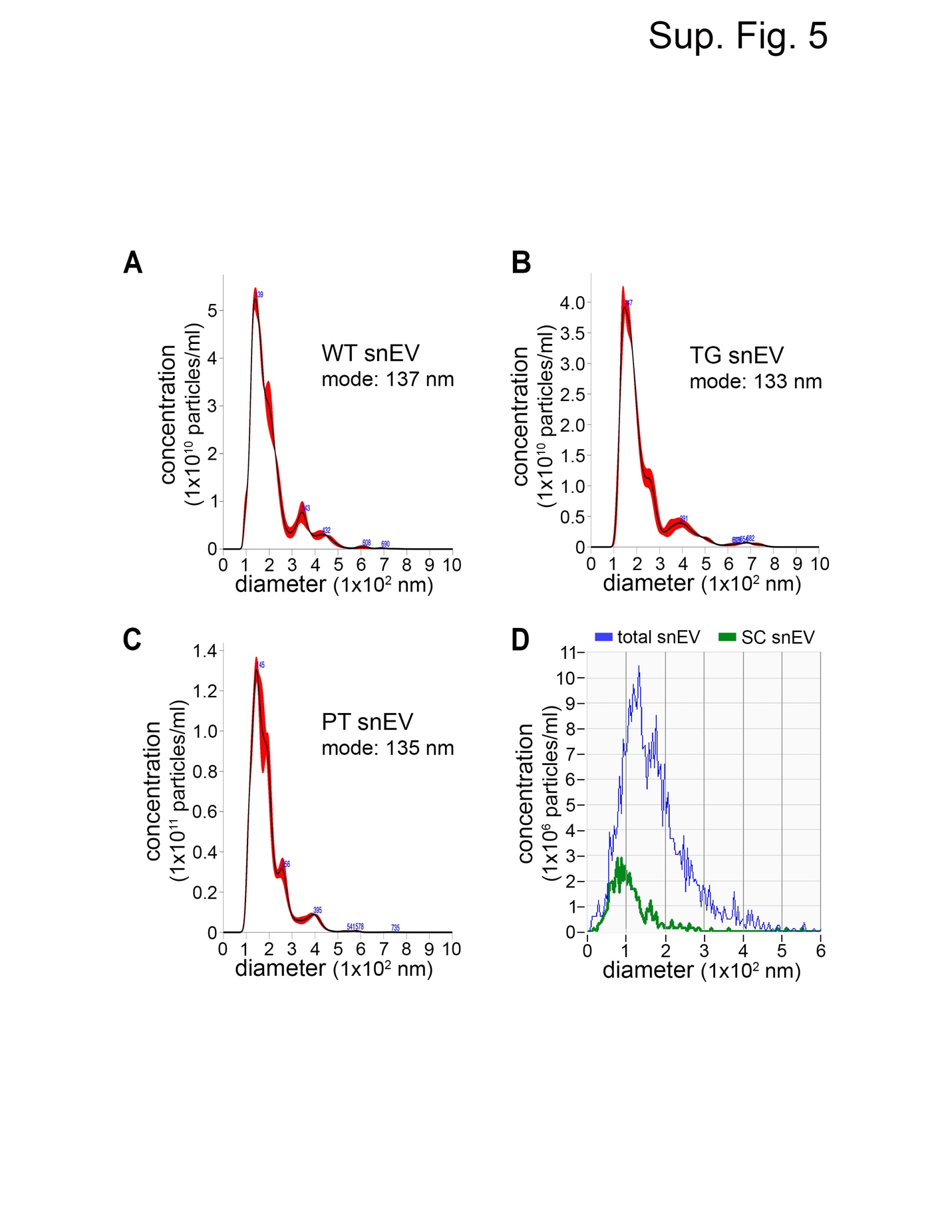

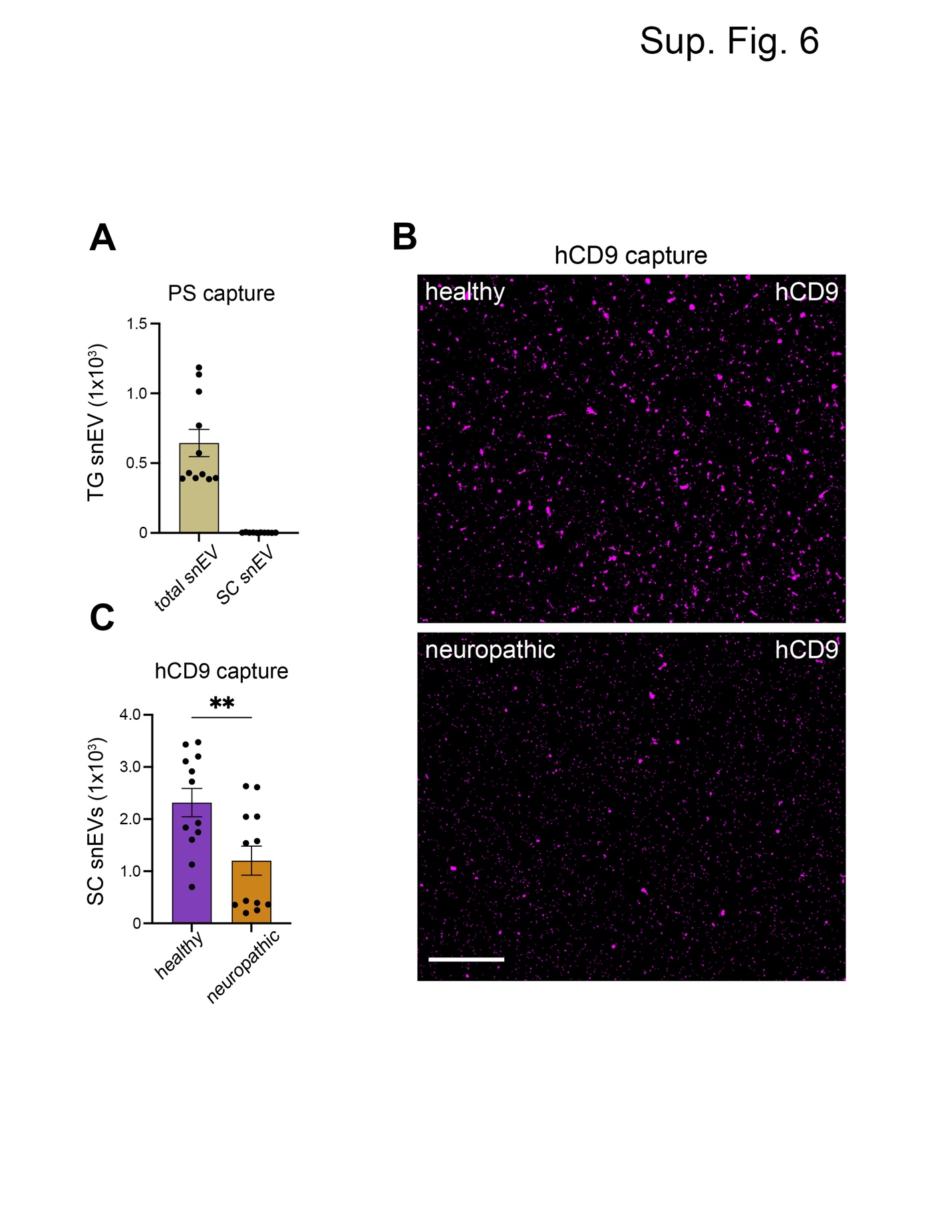

| Table 1. One-fold down regulated differentially expressed proteins (DEP) found in neuropathic snEVs (up regulated in healthy snEV) determined by PEAKS analysis using -10logP with applied false discovery rate control as a global filtering criterion. Uniquely expressed proteins omitted from table. | | |
| --- | --- | --- |
| Protein | **Sig.** | **LFC** |
| Gabarapl2 | 48.07 | 35.44 |
| Prg2 | 89.77 | 34.99 |
| Atp4a | 77.51 | 34.93 |
| Atp1a4 | 77.51 | 34.82 |
| Bche | 104.65 | 34.20 |
| Stmn2 | 69.38 | 34.08 |
| Epx | 75.05 | 33.77 |
| Rps27l | 109.50 | 33.21 |
| Rps27 | 109.50 | 33.21 |
| Dync1li2 | 85.29 | 32.82 |
| Hnrnpab | 74.25 | 32.82 |
| Mtpn | 77.38 | 32.32 |
| Commd6 | 73.03 | 32.00 |
| Calcoco1 | 42.35 | 31.86 |
| Tax1bp3 | 86.40 | 31.84 |
| Dnajb6 | 69.23 | 31.76 |
| Prune1 | 72.29 | 31.66 |
| Cyb5r2 | 70.30 | 31.30 |
| Lhfpl2 | 56.82 | 31.24 |
| Tardbp | 66.98 | 31.18 |
| Srsf3 | 90.88 | 31.09 |
| Pdxp | 66.62 | 31.07 |
| Colgalt1 | 66.66 | 30.65 |
| Mydgf | 49.27 | 30.62 |
| Zbtb8os | 70.26 | 30.56 |
| Rpl35a | 89.44 | 30.47 |
| Lactb2 | 71.68 | 29.91 |
| Ncs1 | 57.58 | 29.81 |
| Pir | 65.52 | 29.76 |
| Gng4 | 16.66 | 2.06 |
| Pvalb | 17.04 | 1.35 |
| Rab1A | 15.99 | 1.30 |
| Cd9 | 20.12 | 1.25 |
| Ahcyl2 | 17.74 | 1.05 |
| Macroh2a2 | 21.34 | 1.05 |
| Rab6a | 15.7 | 0.99 |
| Cd200 | 17.49 | 0.90 |
| Macroh2a1 | 21.34 | 0.80 |
| Fxyd1 | 17.22 | 0.78 |
| Prkcg | 15.06 | 0.74 |
| Prkaa2 | 19.64 | 0.71 |
| Spcs1 | 16.53 | 0.69 |
| Rhoa | 18.28 | 0.65 |
| Ptgis | 19.45 | 0.56 |
| Nme1 | 15.06 | 0.56 |
| Sec23a | 18.15 | 0.55 |
| Prkca | 15.06 | 0.53 |
| Anp32c | 26.15 | 0.47 |
| Bub3 | 16.62 | 0.46 |
| H4f16 | 15.83 | 0.42 |
| Hprt1 | 17.13 | 0.40 |
| Gm7298 | 15.27 | 0.40 |
| Mug1 | 15.27 | 0.36 |
| Ckb | 17.54 | 0.35 |
| Anp32a | 26.15 | 0.35 |
| Mug2 | 15.27 | 0.34 |
| 4930544G11Rik | 18.28 | 0.31 |
| Nde1 | 16.17 | 0.26 |
| Stxbp1 | 16.71 | 0.24 |
| Adam23 | 15.22 | 0.22 |
| Cct7 | 16.63 | 0.19 |
| Gm3839 | 15.61 | 0.14 |
| Cp | 20.71 | 0.14 |
| Rap2a | 16.41 | 0.13 |
| Gapdh | 15.61 | 0.11 |
| Pgk1 | 16.99 | 0.11 |
| Gm10358 | 15.61 | 0.1 |
| Nt5e | 16.5 | 0.07 |
| Rab14 | 16.23 | 0.01 |
| St13 | 15.31 | 0 |

| Table 2. One-fold up regulated differentially expressed proteins (DEP) found in neuropathic snEVs (down regulated in healthy snEVs) determined by PEAKS analysis using -10logP with applied false discovery rate control as a global filtering criterion. Uniquely expressed proteins omitted from table. | | |
| --- | --- | --- |
| Protein | **Sig.** | **LFC** |
| Acot4 | 104.06 | 34.31 |
| Acot6 | 104.06 | 34.31 |
| Acot5 | 104.06 | 34.31 |
| Acot1 | 104.06 | 34.30 |
| Acot2 | 104.06 | 34.30 |
| Krt80 | 66.30 | 34.00 |
| Acat3 | 86.04 | 33.77 |
| Acat2 | 86.04 | 33.74 |
| Ide | 64.05 | 33.25 |
| EphB3 | 93.39 | 33.11 |
| Erg28 | 71.97 | 32.46 |
| Rpl35 | 56.36 | 32.06 |
| Efhd2 | 51.39 | 29.32 |
| Fa2h | 19.42 | 5.25 |
| Fgf1 | 16.78 | 4.55 |
| Pcnp | 18.52 | 1.74 |
| Anp32b | 26.15 | 1.57 |
| Snx12 | 15.61 | 1.54 |
| Ago2 | 15.77 | 1.25 |
| Rpl14 | 18.19 | 1.20 |
| Ehbp1 | 17.38 | 1.12 |
| Dnajc5 | 27.93 | 1.04 |
| Agpat1 | 36.38 | 1.04 |
| Nradd | 16.01 | 1.02 |
| Ubl3 | 15.56 | 1.00 |
| Fkbp1b | 37.65 | 0.97 |
| Emilin3 | 22.81 | 0.82 |
| Nptn | 27.65 | 0.69 |
| Septin10 | 17.4 | 0.63 |
| Hars2 | 15.75 | 0.55 |
| Myo5a | 17.43 | 0.53 |
| Rabep1 | 21.78 | 0.49 |
| Gapdhs | 15.61 | 0.49 |
| Zyx | 17.67 | 0.49 |
| Ehd3 | 16.32 | 0.41 |
| Pcyox1l | 18.52 | 0.35 |
| Xpnpep2 | 22.77 | 0.34 |
| Hars1 | 15.75 | 0.31 |
| Il6st | 18.55 | 0.30 |
| Glg1 | 18.18 | 0.29 |
| Crmp1 | 19.32 | 0.28 |
| Dip2b | 19.25 | 0.26 |
| Pgk2 | 16.99 | 0.23 |
| Susd2 | 16.49 | 0.09 |
| Npr2 | 28.59 | 0.09 |
| Arpc1b | 19.46 | 0.06 |
| Prkaa1 | 19.64 | 0.01 |
